# Large-scale single-cell long-read genomics enables high-resolution microbiome profiling

**DOI:** 10.1101/2024.09.10.612220

**Authors:** Mengzhen Li, Shiyan Li, Hengxin Liu, Xuanpei Zhai, Jie Li, Yanan Du, Xiaolu Li, Jiawei Tong, Jian Zhang, Rong Zhang, Zhangyue Song, Quanjiang Ji, Yuan Luo, Ting Zhang, Wu Wei, Yifan Liu

## Abstract

Microbial communities are extraordinarily diverse and play crucial roles in health and disease, yet current methods lack the resolution and scalability needed to dissect their genomic and ecological complexity at the single-cell level. Here, we present CAP-seq, a high-throughput single-microbe genomics platform that combines hydrogel-based semi-permeable encapsulation with minimal microfluidics to recover thousands of single-amplified genomes (SAGs) with long reads and high completeness at low sequencing depth. We benchmarked CAP-seq using defined microbial communities, demonstrating strain-level resolution, accurate detection of rare taxa, and genome recovery exceeding 50% at ∼10× coverage. Applying CAP-seq to pediatric *Clostridioides difficile* infection microbiomes, we generated a high-resolution single-cell atlas comprising tens of thousands of SAGs across hundreds of species. Host-resolved profiling of the cryptic plasmid pBI143 revealed previously hidden low-abundance host associations, six new plasmid versions, and their coexistence within individuals, indicating complex plasmid evolution in situ. Longitudinal analysis during fecal microbiota transplantation and vancomycin treatment uncovered dynamic remodeling of microbial hosts, antimicrobial resistance genes, and plasmids at single-cell resolution. CAP-seq enables scalable, high-performance single-cell genomics and provides a practical, widely accessible platform for microbiome analysis, paving the way for large-scale exploration of microbial dark matter and host–microbe interactions across diverse ecosystems.

## INTRODUCTION

Microorganisms are the most abundant and diverse living forms on Earth^1,2^. Current estimates suggest that up to a trillion microbial species exist^2–4^, distributed across natural environments and the human body. Yet, microorganisms in most ecosystems remain under-characterized, representing a major component of “microbial dark matter”^5^. Even within the human microbiome, our understanding of community structure and function remains limited despite notable efforts such as the Human Microbiome Project^6^. Resolving these complex biological systems ultimately requires the ability to define the identity and function of each microbial strain, or even individual cells.

Genomic information is a fundamental determinant of microbial identity and function. Traditionally, microbial genomes are obtained either by culturing and sequencing isolated strains^7^ or by metagenomic assembly^8,9^. Both strategies have intrinsic limitations for studying complex communities: culturing is constrained by the cultivability and throughput of strains, whereas metagenomics lacks single-cell and strain-level resolution. Recent advances in high-throughput biotechnology have enabled single-cell microbial genomics by isolating cells using microfluidics followed by individual genome amplification and indexing^10,11^. However, existing methods recover only a limited fraction of genomes and require multiple serial microfluidic operations, limiting scalability and accessibility. There remains a pressing need for high-coverage, practical approaches that can be broadly applied to diverse microbiomes.

Here, we present CAP-seq, a scalable and efficient method for recovering thousands of high-quality single-amplified genomes (SAGs) by processing individual cells in compartments with adjusted permeability (CAPs). CAPs are micrometer-scale hydrogel capsules that retain intracellular DNA while allowing diffusion of reagents and buffers^12,13^. This semi-permeable structure enables lysis, whole-genome amplification, tagmentation, and washing to be performed in bulk, improving reaction efficiency and data quality (Scheme 1). Importantly, CAP-seq requires only two microfluidic steps: encapsulation and barcoding (Fig. 1a and Scheme 1), which significantly reduces operational complexity compared with current state-of-the-art approaches.

**Figure 1.**
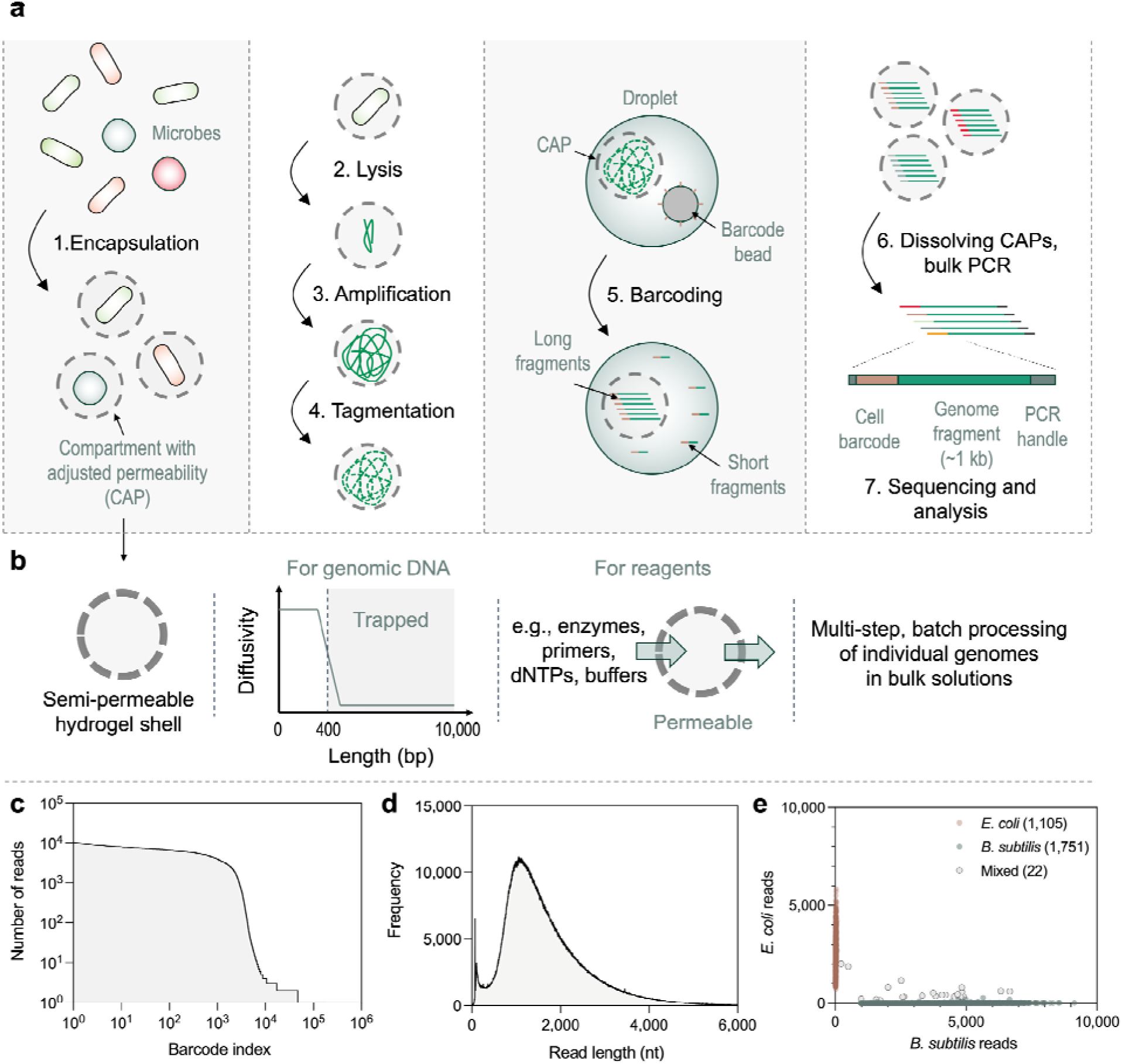
Schematic and preliminary evaluation of CAP-seq. (a) Schematic of single-cell genomic library preparation with CAP-seq. The workflow encapsulates and processes single microbial cells in hydrogel compartments with adjusted permeability (CAPs), generating amplified and barcoded single-cell genomes suitable for long-read sequencing. CAP-seq involves two microfluidic steps, highlighted with a grey background: (1) cell encapsulation and (2) genome barcoding. (b) A key feature of CAP-seq – the use of hydrogel capsules to compartmentalize cells. The capsules are semi-permeable to dsDNA with a cutoff size of ∼400 bp, therefore retaining cell genomes and large genomic DNA fragments. They remain permeable to all other reagents of CAP-seq, including enzymes, barcode primers, dNTPs, and buffers, thus enabling multi-step batch processing of individual genomes in bulk solutions. (c&d) Evaluation of CAP-seq using a 1:1 mixture of *Escherichia coli* and *Bacillus subtilis* cells. (c) Read distribution across different barcode groups (or single-amplified genomes, SAGs). (d) Read length profile after barcode trimming. Bin size: 1. (e) Scatter plot depicting the number of *E. coli* and *B. subtilis* reads associated with SAG.

We critically evaluated CAP-seq using defined microbial communities. A single experiment yielded thousands of high-purity SAGs, achieving >50% genome coverage at ∼10× sequencing depth. Long-read sequencing produced kilobase-level barcoded reads, enabling strain-level resolution and recovery of mobile genetic elements such as plasmids and phages. We then applied CAP-seq to profile the gut microbiomes of pediatric patients with *Clostridioides difficile* infection (CDI)^14^, a clinically relevant model of dysbiosis^15^. CAP-seq generated a single-cell atlas comprising tens of thousands of SAGs across hundreds of species, revealing both shared and patient-specific microbial lineages. Host-resolved plasmid profiling uncovered extensive ecological and genomic diversity of the cryptic plasmid pBI143^16^, including previously undetected low-abundance host associations and four novel plasmid versions that frequently coexisted within individual patients. Finally, longitudinal analyses during fecal microbiota transplantation^17^ (FMT) and vancomycin treatment revealed dynamic shifts in microbial hosts, antimicrobial resistance genes, and plasmids, reflecting complex ecological restructuring during therapy. Together, these results establish CAP-seq as a powerful and practical platform for large-scale, high-resolution single-cell genomics, enabling new insights into the structure, dynamics, and evolution of complex microbial communities.

## RESULTS

### CAP-seq workflow

The CAP-seq workflow begins by encapsulating individual microbial cells within the CAPs (Fig. 1a). This is achieved by co-flowing a cell suspension and hydrogel precursor solutions through a microfluidic droplet generator (Supplementary Fig. 1a). During emulsification, the precursors, dextran and acrylate-modified poly(ethylene glycol) diacrylate (PEGDA), undergo liquid–liquid phase separation^18^, forming ∼60 µm microcapsules that encapsulate single cells within the inner dextran-rich core^19^. The PEGDA shell acts as a semi-permeable barrier with an effective cutoff of ∼400 bp for double-stranded DNA (dsDNA). We confirmed this property using a PCR-based permeability assay we developed previously^12,13^ (Supplementary Figs. 1c–e). In addition to dsDNA, the capsule walls remain permeable to all reagents required for subsequent steps, including enzymes, primers, dNTPs, and buffers (Fig. 1b and Scheme 1).

Following encapsulation, cells are lysed in two sequential steps to ensure efficient genomic DNA release. First, the CAPs are incubated in a lysozyme-containing cocktail to digest bacterial cell walls. This is followed by extensive washing and incubation in a second cocktail containing proteinase K, which degrades proteins and completes cell lysis. The lysed CAPs are then transferred to a multiple displacement amplification (MDA) mixture for whole-genome amplification. Subsequently, amplified genomes are tagmented within the capsules using Tn5 transposase.

After amplification and tagmentation, the CAPs are re-encapsulated in droplets and paired with DNA barcode beads for genome indexing. This is performed using a custom-designed droplet merging device^20^ (Supplementary Fig. 1b and Movie 1). Upon merging, the beads dissolve and release unique barcode primers, which diffuse into the CAPs and label the tagmented DNA by linear PCR, thereby assigning each genome a unique cell barcode. The barcoded CAPs are then recovered from the emulsion, washed, and dissolved to release barcoded DNA fragments, which are further amplified in bulk PCR. Finally, sequencing libraries are prepared using standard Oxford Nanopore protocols. The complete CAP-seq workflow is illustrated in Scheme 1 and described in detail in Methods.

CAP-seq of a 1:1 mixture of *E. coli* and *B. subtilis*.

We first characterized CAP-seq using a 1:1 mixture of *E. coli* and *B. subtilis* cells. We performed CAP-seq on around 10,000 cells and obtained 17.0 million raw reads. Our in-house quality control and binning pipeline identified 10.8 million reads with full-length and correct cell barcodes. We binned the reads based on the cell barcodes and obtained 2,878 barcode groups of ≥1,000 reads, including 1,560 groups with ≥3,000 reads (Fig. 1c). After barcode trimming, the remaining reads (genome fragments) mainly distributed from ∼500 to 3,000 nt, with an average value of 1,702 nt (Fig. 1d). We compared the 2,878 barcode groups (≥1,000 reads) to the reference genomes, and successfully assigned 1,105 (38.4%) barcode groups to *E. coli* (98.8% reads on average mapped to the *E. coli* genome) and 1,751 (60.8%) to *B. subtilis* (average of 99.5% mapped reads), Fig. 1e. Only a small fraction (0.8%) of groups were identified as mixed populations. These results confirm the single-cell resolution of CAP-seq and verify that the CAPs effectively preserved the encapsulated single-cell genomes and their amplicons. We next calculated the genome coverage of each barcode group based on read mapping and found that the coverages had not yet reached saturation for either species (Supplementary Fig. 1f). Nonetheless, CAP-seq achieved reasonable coverage performance. The *E. coli* groups (N = 1,105) had a mean coverage of 14.2% at a mean sequencing depth of 0.73×, while the *B. subtilis* groups (N = 1,751) exhibited a mean coverage of 44.8% at a mean sequencing depth of 1.39×.

### CAP-seq of a mock microbial community

To further evaluate CAP-seq, we sequenced a mock microbial community composed of four bacterial species with varied Gram polarities at uneven ratios (Fig. 2a). We sequenced about 10,000 cells and obtained 88 million raw reads. We then obtained 3,490 barcode groups containing over 1,000 reads (52 million reads with average read length 1,318 after barcode trimming, Supplementary Figs. 2a&b), 90% of which had a purity (percent reads mapped to a dominant genome) of over 95%, and 83% with over 99% purity (Fig. 2b). Notably, CAP-seq consistently captured rare species across independent experimental replicates, demonstrating its high sensitivity and reproducibility benefiting from its intrinsic single-cell resolution (Fig. 2c). After mapping to reference genomes, we assessed the completeness of the 3,490 SAGs (Supplementary Fig. 2c). Because genome completeness depends on sequencing depth, we plotted coverage as a function of depth (Fig. 2d). It can be found that most SAGs reached >50% coverage at only ∼10× sequencing depth, independent of Gram status, and coverage further increased to ∼80% at around 20X depth. To further evaluate SAG quality in settings where no reference genomes are available, we performed *de novo* assembly of the 3,490 SAGs and assessed the obtained assemblies with a marker gene-based, reference-free tool (CheckM). These *de novo* assemblies yielded completeness up to ∼60% (mean 26.61%; Fig. 2e), average contamination of 1.48% (Supplementary Fig. 2d) and N50 of 14,370 (Fig. 2f), at an average sequencing depth of 2.69× per SAG. These results highlight CAP-seq’s ability to yield high-quality SAGs with moderate sequencing effort.

**Figure 2.**
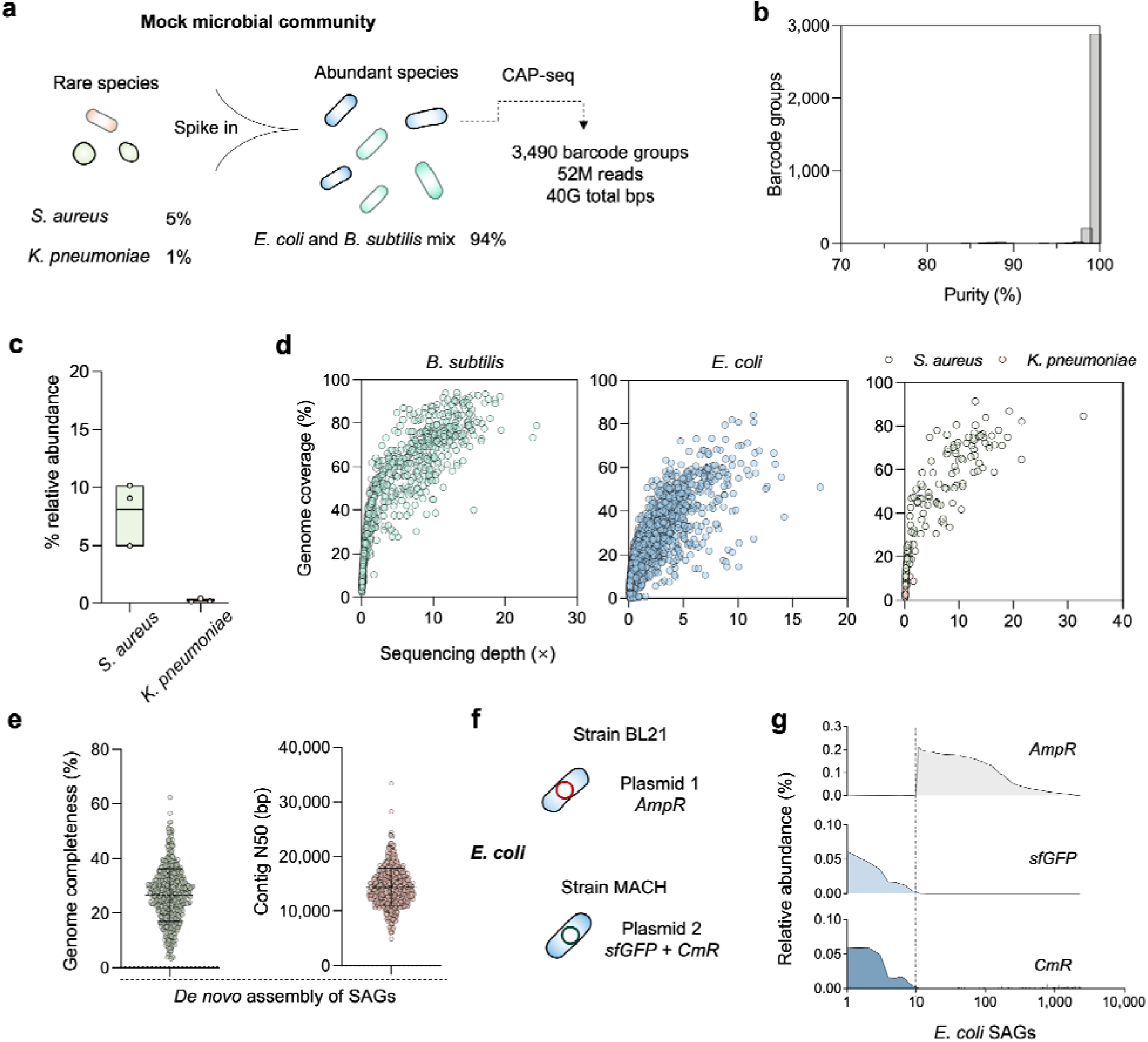
Evaluation of CAP-seq using a mock microbial community. (a) Schematic of the mock community experiment. A mixture containing abundant species (*E. coli, B. subtilis*) and spiked rare species (*Staphylococcus aureus, Klebsiella pneumoniae*) was processed by CAP-seq, yielding 3,940 barcode groups and 97 Gb total bases. (b) Purity distribution of all SAGs obtained from the mock community, showing that 90% of SAGs exceeded 95% purity (83% SAGs over 99% purity). (c) Relative species abundance profiles inferred from CAP-seq. The results of two more parallel experiments were shown to evaluate the reproducibility of CAP-seq in detecting rare bacterial species (N = 3). (d) Genome coverage versus sequencing depth for SAGs from *B. subtilis* (N = 877), *E. coli* (N = 2,427), *S. aureus* (N = 173), and *K. pneumoniae* (N = 13). (e) Quality metrics of *de novo* assemblies from 3,490 SAGs, showing completeness (left) and contiguity (N50; right). The completeness was evaluated using a reference-free, marker gene-based tool (CheckM). (f) Experimental design for distinguishing *E. coli* strains BL21 and MACH1 based on plasmid-encoded markers (*AmpR* vs. *sfGFP* + *CmR*). The two strains were mixed at a ∼100:1 ratio (BL21:MACH1). (g) Relative abundance of *AmpR*, *sfGFP*, and *CmR* genes across 2,427 *E. coli* SAGs, demonstrating strain-level resolution and marker specificity. The SAGs are sorted in descending order of *sfGFP* and then *AmpR*.

We further analyzed the *E. coli* groups, which consisted of two distinct strains, BL21 and MACH1 (Fig. 2f). These strains were engineered to harbor strain-specific plasmid-encoded markers: BL21 carried *AmpR*, whereas MACH1 carried *sfGFP* and *CmR*. Gene profiling across 2,427 *E. coli* SAGs revealed distinct relative abundance patterns (Fig. 2g). The abundances of *sfGFP* and *CmR* were highly correlated and predominated in SAGs lacking *AmpR*. In contrast, the remaining SAGs contained only *AmpR* without *sfGFP* or *CmR*. These distinct marker patterns enabled unambiguous strain-level separation, assigning 10 SAGs to MACH1 and the remainder to BL21. Collectively, these results demonstrate the capacity of CAP-seq to resolve intraspecific heterogeneity at single-cell resolution.

### Single-cell profiling of gut microbiota in pediatric CDI patients

After validating the performance of CAP-seq in mock communities, we applied the method to clinical samples. We focused on *Clostridioides difficile* infection, a challenging disease often associated with antibiotic resistance and typically occurring in patients with prior antibiotic exposure^21,22^. We recruited four pediatric CDI patients and divided them into two treatment groups (Fig. 3a): two received vancomycin therapy (VAN-1 and VAN-2) and two underwent fecal microbiota transplantation (FMT-1 and FMT-2). All patients had a history of antibiotic exposure prior to CDI diagnosis. The stool samples were collected at two to three time points: at diagnosis (or pre-treatment), 2 weeks after (the end of 2-week vancomycin treatment), and, when available, 4 weeks after the diagnosis. In total, 10 stool samples were processed using CAP-seq, yielding ∼160 Gb of long-read data per sample (average read length per sample ranged from 962 to 2009; Supplementary Table 1). After quality control and barcode binning, each sample produced between 1,646 and 16,820 SAGs (Supplementary Fig. 3a). *De novo* assembly followed by taxonomic annotation assigned ∼76.3% of SAGs to species level, with each sample containing 32–152 species (Supplementary Fig. 3b). When mapping SAGs to reference genomes, we observed that genome coverage from clinical stool samples was generally lower than that obtained from mock samples, with most SAGs not exceeding 30% coverage (Supplementary Fig. 3c). While lower sequencing depths partly explain this reduction, additional factors such as cell damage introduced during sampling, transportation, storage, and purification of clinical stool samples are also likely to contribute.

**Figure 3.**
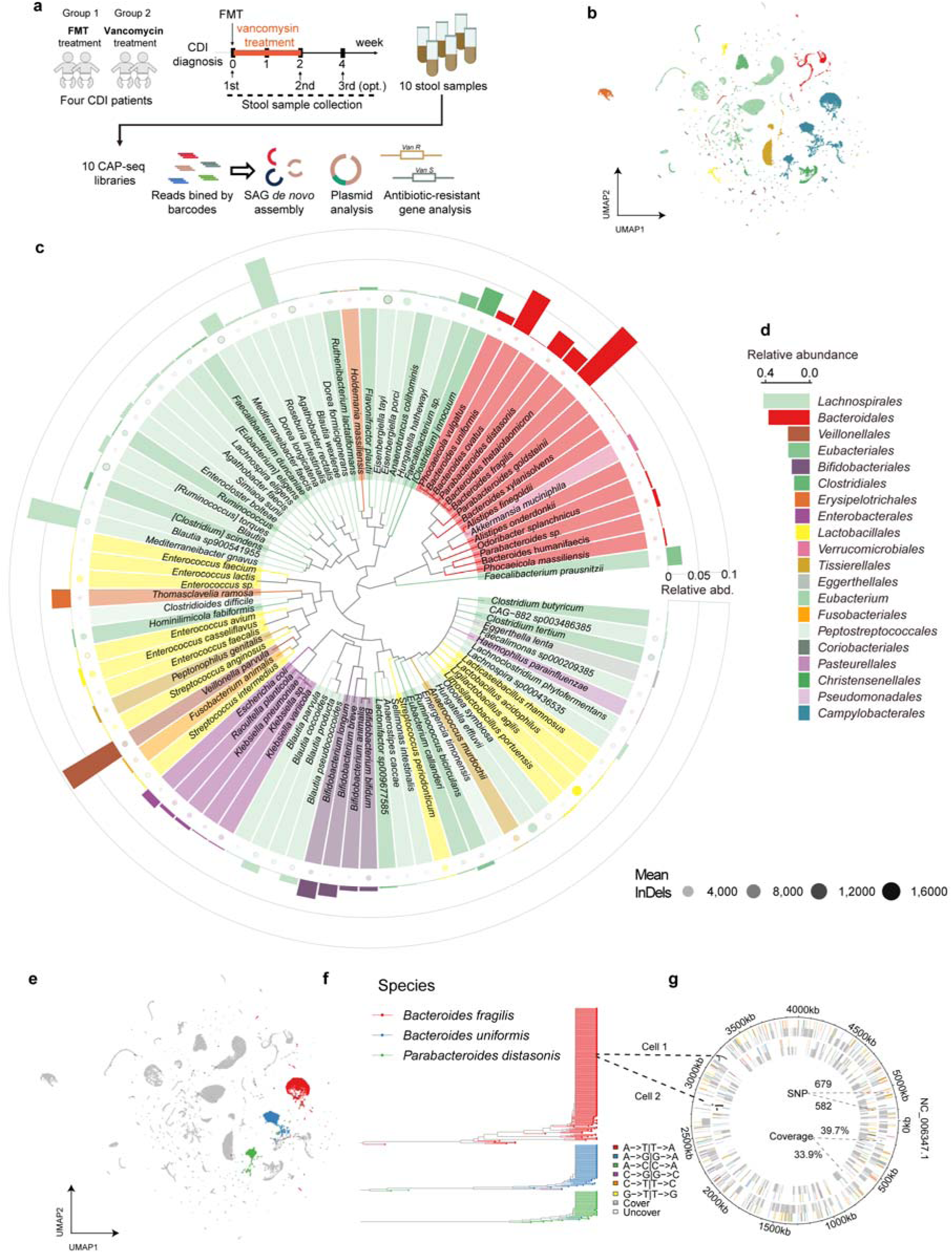
Single-cell atlas of gut microbial communities in *Clostridioides difficile* infection (CDI) patients during treatment. (a) Schematic of the study design. Stool samples were collected longitudinally from four CDI patients receiving either Fecal microbiota transplant (FMT; patients FMT-1 and FMT-2) or vancomycin treatment (patients VAN-1 and VAN-2), at pre-treatment, week 2, and optional week 4 time points. A total of 10 CAP-seq libraries were generated and analyzed for microbial genomes, plasmids, and antimicrobial resistance genes. (b) Uniform Manifold Approximation and Projection (UMAP) for of 54,532 single-amplified genomes (SAGs) from all samples, colored by taxonomic order, illustrating the microbial diversity captured by CAP-seq. (c) Phylogenetic tree and relative abundance profiles of the top 100 recovered microbial species. Bar lengths indicate relative abundances; bar colors correspond to taxonomic orders. Dot sizes represent mean numbers of detected InDels per species. (d) Order-level composition of the CDI gut microbiome, showing the relative abundances of major bacterial orders recovered by CAP-seq across all samples. Colors correspond to taxonomic orders and match panels b and c. (e) UMAP visualization highlighting SAGs assigned to *Bacteroides fragilis* (red), *Bacteroides uniformis* (blue), and *Parabacteroides distasonis* (green). (f) Single-cell phylogenies of these three species, revealing within-species genomic diversity and population structure across samples. (g) Example of single-cell–resolved single-nucleotide polymorphism (SNP) detection: two *B. fragilis* cells from the same sample exhibit hundreds of distinct SNPs across their genomes, illustrating CAP-seq’s ability to resolve intra-species genetic variation at single-cell resolution.

To obtain a global view of the CDI microbiomes, we first combined all 10 samples (∼54,000 SAGs and 326 bacterial species) and constructed an ANI-based similarity matrix by mapping each SAG to all reference genomes. Uniform manifold approximation and projection (UMAP) visualization of this matrix revealed distinct clusters corresponding to different bacterial species, thus generating a single-cell atlas of the pediatric CDI microbiomes (Fig. 3b). Importantly, the maximum ANI within each cluster exceeded 95% (Supplementary Figs. 3d&e), supporting the accuracy of species-level assignments and the high single-cell purity of CAP-seq. The UMAP projection captured microbial lineages shared across individuals, with patient-specific clustering patterns also evident in sample-wise analyses (Supplementary Fig. 3f). To further characterize community composition, we ranked the species by relative abundance across the combined dataset. Fig. 3c shows the top 100 species, together with their relative abundances across the 10 samples. The community was dominated by members of the orders *Bacteroidales* and *Lachnospirales* (Fig. 3d), but also included taxa relevant to CDI pathogenesis and dysbiosis, such as *C. difficile*, *Enterococcus*, and *Klebsiella*. Specifically, *C. difficile* was detected in three of the four patients. However, its relative abundance was consistently low, with all samples showing <1% of SAGs assigned to *C. difficile* (Supplementary Fig. 3g). This observation is consistent with prior microbiome studies of CDI, which also reported that *C. difficile* typically remains a minority member of the gut community despite its disproportionate role in disease pathogenesis^23,24^.

In addition to species abundance, CAP-seq enabled quantification of insertion–deletion (InDel) events at the single-cell level. The average number of InDels per species is shown by the size of the outer circles in Fig. 3c, ranging from a few hundred to over 8,000 events. Notably, certain taxa such as *Bacteroides fragilis* and *Parabacteroides distasonis* exhibited both high relative abundance and elevated mean InDel counts, suggesting active genomic plasticity in these lineages. In contrast, low-abundance taxa such as *Enterococcus spp.* carried relatively fewer InDels, consistent with more conserved genomes.

To further resolve species-level population structures within the CDI microbiomes, we focused on the dominant *Bacteroides* lineages. UMAP projection of SAGs highlighted distinct clusters corresponding to *B. fragilis*, *Bacteroides uniformis*, and *P. distasonis* (Fig. 3e). These clusters were consistent with species-level taxonomy, confirming the ability of CAP-seq to capture population structure at single-cell resolution. We then constructed a phylogenetic tree based on single-nucleotide polymorphisms (SNPs) across the SAGs, which further delineated the three *Bacteroides* species into well-separated clades (Fig. 3f). The tree topology supported both interspecies divergence and intraspecies microdiversity, revealing sub-lineages within each cluster that likely reflect strain-level variation. As a representative example, we highlighted two *B. fragilis* SAGs from the tree (Fig. 3g). Mapping reads from these SAGs to the reference genome yielded 33.9% and 39.7% coverage, and identified 582 and 679 SNPs distributed across the genome, respectively. The diversity of substitution types (e.g., A→T, A→C, G→C) illustrates the fine-scale genomic heterogeneity present even within a single species. Together, these results underscore CAP-seq’s ability to link phylogenetic microdiversity with concrete genomic variation at single-cell resolution.

### Single-cell host-resolved profiling of plasmid pBI143 within CDI stool microbiomes

The single-cell resolution of CAP-seq enables direct profiling of mobile genetic elements, such as plasmids and phages, within individual microbial cells. Here, we focused on the cryptic plasmid pBI143 (Fig. 4a), which was recently reported as one of the most abundant genetic elements in the human gut microbiome, surpassing even phages in industrialized populations, and has been associated with inflammatory bowel disease^16^ (IBD). We first searched for pBI143 in CDI stool microbiomes and detected it in two patient samples (FMT-2 and VAN-2), where it was present in approximately 39% of all SAGs (Fig. 4b). Single-cell quantification of plasmid copy number revealed a broad dynamic range, with two major peaks around 1 and 100 copies per cell and maximum values exceeding 10 copies (Fig. 4c). Such bimodal distributions likely reflect alternative replication states across different host backgrounds or ecological contexts, with low-copy maintenance supporting stable vertical transmission and high-copy amplification occurring under specific physiological or ecological conditions^25,26^.

**Figure 4.**
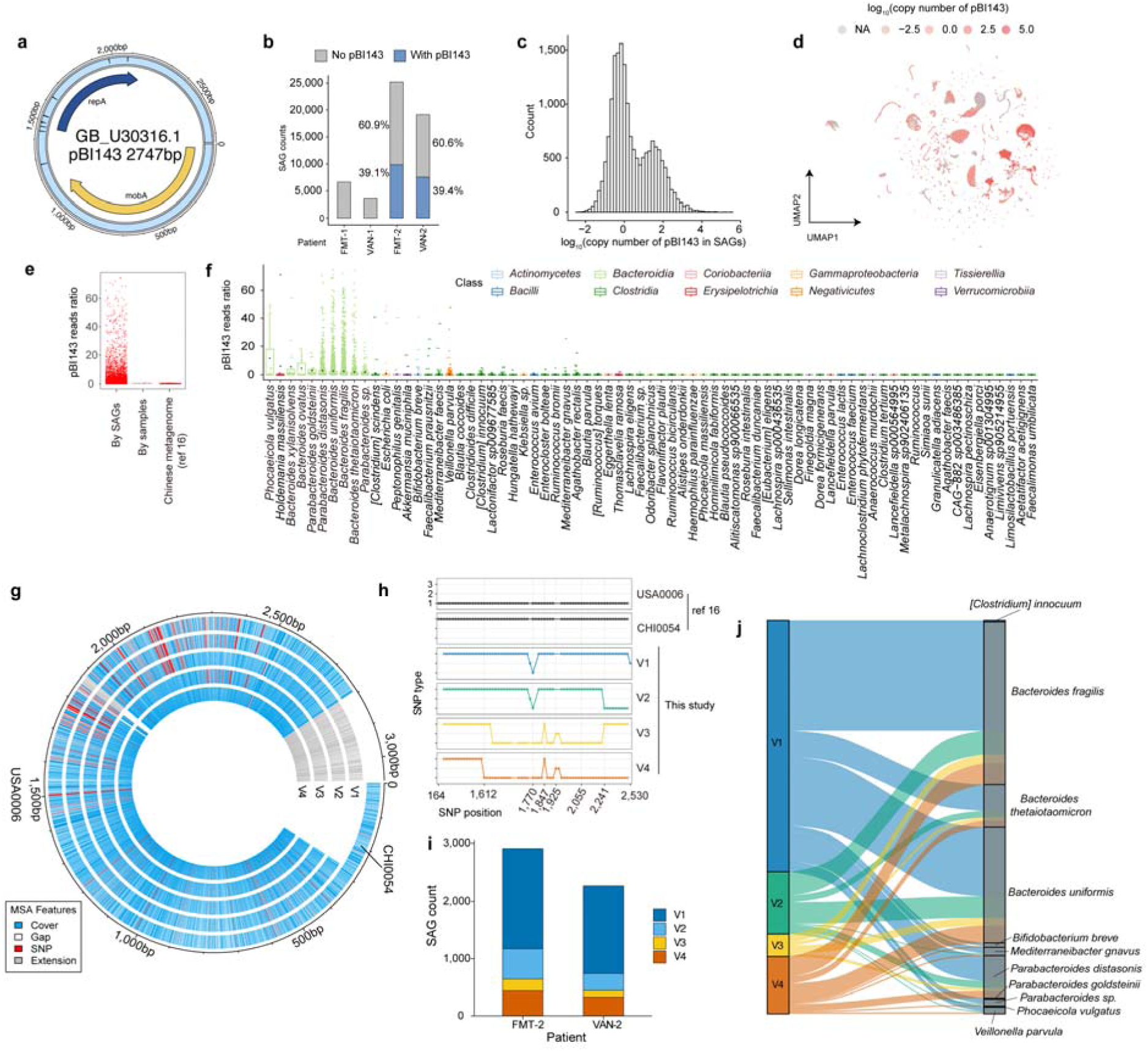
Single-cell profiling reveals extensive diversity and ecological distribution of plasmid pBI143 in CDI gut microbiomes. (a) Genome organization of pBI143 (GeneBank_U30316.1, 2747 bp). The inner ring shows the aggregate read coverage across all SAGs mapped to the plasmid, illustrating overall recovery across the entire sequence. (b) Proportion of SAGs containing pBI143 across four CDI patient samples. (c) Distribution of pBI143 copy number per cell. (d) UMAP projection of all SAGs colored by plasmid copy number, illustrating heterogeneity in plasmid carriage across microbial clusters. (e) Ratio of plasmid-positive SAGs and by-sample abundance for each species, compared with the Chinese metagenomic data from reference 16. (f) Species-level distribution of plasmid-positive cells. (g) Comparative genomics of pBI143 versions. *De novo*–assembled plasmid-covering contigs were clustered based on SNP profiles, revealing four novel versions (V1–V4) distinct from previously described USA0006 and CHI0054 lineages. (h) SNP classification relative to USA0006 and CHI0054 references, showing that V1 closely resembles CHI0054, whereas V2–V4 exhibit mosaic SNP patterns. (i) Abundance of plasmid versions across patients, illustrating that multiple versions coexisted within individual hosts. (j) Host range of the four plasmid versions.

Projecting plasmid copy number onto the single-cell microbiome atlas revealed its presence across 73 microbial clusters, with substantial heterogeneity among host species (Fig. 4d). We then questioned why pBI143 was not detected in FMT-1 and VAN-1. One possibility was that these patients lacked key host species typically associated with pBI143 carriage. However, major host taxa with high plasmid copy numbers (>20) identified in FMT-2 and VAN-2 were also present in FMT-1 and VAN-1 (Supplementary Fig. 4a), indicating that the absence of pBI143 is not simply due to missing host species but likely reflects ecological or evolutionary differences in plasmid colonization across individuals.

We next examined how plasmid abundance varies across single cells. A subset of SAGs carried unusually high plasmid loads, with pBI143 reads accounting for more than 20% of total reads in some cells (Fig. 4e). When analyzed at the bulk sample level by aggregating SAG reads, pBI143 accounted for less than 1% of total sequences (0.46%), consistent with previous metagenomic analyses of Chinese gut microbiomes^16^. This highlights the unique advantage of single-cell sequencing in revealing high-copy plasmid carriage that would otherwise be masked in bulk analyses.

We further explored the host distribution of pBI143 across all SAGs (Fig. 4f). Previous metagenomic studies reported that pBI143 is largely confined to members of the class *Bacteroidia*, mainly within the genera *Bacteroides*, *Phocaeicola*, and *Parabacteroides*^16^. Consistent with previous work, our data showed that *Bacteroidia* taxa indeed exhibited high plasmid abundance. However, CAP-seq further revealed additional host lineages beyond this class, such as *Gammaproteobacteria*, *Bacilli*, and *Clostridia*, which had not been previously linked to pBI143. These newly identified hosts generally have a low abundance of pBI143, which may explain why they were overlooked in bulk metagenomic analyses. This expanded host spectrum demonstrates that pBI143 is not limited to a single phylogenetic group but can circulate across multiple bacterial classes within the gut ecosystem. By directly linking plasmids to their cellular hosts, CAP-seq overcomes the resolution barrier of metagenomics and uncovers hidden plasmid–host associations that are otherwise inaccessible in bulk community sequencing.

To further dissect host–plasmid interactions, we examined the fraction of plasmid-positive cells within each species (Supplementary Fig. 4b). Species with the highest carrier fractions were not always those with the highest plasmid abundance. For example, *Lancefieldella sp000564995* exhibited >75% pBI143-positive cells despite low per-cell plasmid abundance (average 2.6%), suggesting widespread but low-copy maintenance strategies. Conversely, *Phocaeicola vulgatus* displayed lower carrier fractions but higher plasmid loads in positive cells, consistent with episodic amplification or variable copy-number control^27^.

Previous metagenomic analyses reported that pBI143 typically exists as a limited number of sequence variants within individual hosts, usually dominated by a single lineage^16^. pBI143 has three previously identified versions, CHI0054, USA0006, and ISR0084, across global populations, with most hosts carrying only one version^16^. However, as bulk metagenomic assemblies average signals across many cells, they may overlook finer-scale genomic diversity within individual hosts. We therefore leveraged CAP-seq to investigate the genomic diversity of pBI143 at single-cell resolution. *De novo* assembly of individual SAGs enabled the recovery of 5,247 contigs covering the full plasmid length. We selected 2,010 high-quality contigs for version typing. By clustering based on SNP profiles, we discovered four plasmid versions (V1– V4) (Supplementary Fig. 4c). Comparative genomic analyses showed that these versions are genetically distinct from CHI0054 and USA0006, exhibiting varying degrees of similarity (Fig. 4g and Supplementary Fig. 4d). The previously described ISR0084 was not included in the main comparison because of its substantial divergence from the pBI143 consensus used here. To further dissect the relationships among plasmid versions, we classified SNPs into USA0006-specific (type 1), CHI0054-specific (type 3), and novel (type 2) categories (Fig. 4h). V1 is most similar to CHI0054, whereas V2, V3, and V4 exhibit mosaic SNP patterns to varying degrees. These observations suggest that these newly identified plasmid versions likely originated through recombination between existing lineages within the gut.

Beyond identifying new pBI143 lineages, our single-cell data show that pBI143 is not monoclonal within individuals. Multiple plasmid versions (V1–V4) coexisted in each patient and often within the same host species (notably *B. fragilis*, *Bacteroides thetaiotaomicron*, and *B. uniformis*), Fig. 4i. This contrasts with the earlier report of host-level monoclonality^16^. Across patients, V1 was the most common, and V3 was rare. We next analyzed version–host associations. All four versions (V1–V4) were detected across multiple species, dominated by the taxa of *Bacteroides* and *Parabacteroides*, in both patients, with differences primarily in relative prevalence rather than host exclusivity (Fig. 4j and Supplementary Fig. 4e). These patterns may suggest that version-specific compatibility and maintenance constraints exist across host backgrounds, rather than strict host restriction. The coexistence of multiple versions within individuals and species, together with their distinct host ranges, points to a dynamic intra-host plasmid population shaped by recombination, horizontal transfer, and competitive maintenance. This revises the prevailing view of pBI143 ecology and reveals previously hidden diversity and population structure in the human gut. These findings also highlight the power of CAP-seq to resolve fine-scale mobile element dynamics at single-cell resolution, enabling direct tracking of plasmid diversification and host association within complex microbial communities.

### Single-cell profiling of treatment-induced shifts in microbial communities and resistance gene dynamics

Having established the static ecological landscape of microbial hosts and their associated plasmids, we moved to investigate how the gut microbiome dynamically responds to therapeutic perturbations during FMT and vancomycin treatment. To begin, we examined species-level compositional changes across treatment stages by visualizing all SAGs over time using UMAP projections (Fig. 5a). Both FMT and vancomycin treatment caused marked community shifts from pre-treatment to week 2 and week 4. It can be noted that these shifts were greater under vancomycin therapy, consistent with the broad-spectrum bactericidal activity of the drug and its strong selective pressure on gut taxa. We next used pairwise Jaccard similarity to quantify compositional overlap between consecutive sampling stages (Supplementary Fig. 5a). Consistent with the visual pattern, FMT-treated microbiomes showed significantly higher Jaccard scores (mean 0.41) than those receiving vancomycin (mean 0.10), indicating greater temporal stability and less turnover of species composition. This finding supports that FMT promotes community restoration rather than wholesale restructuring.

**Figure 5.**
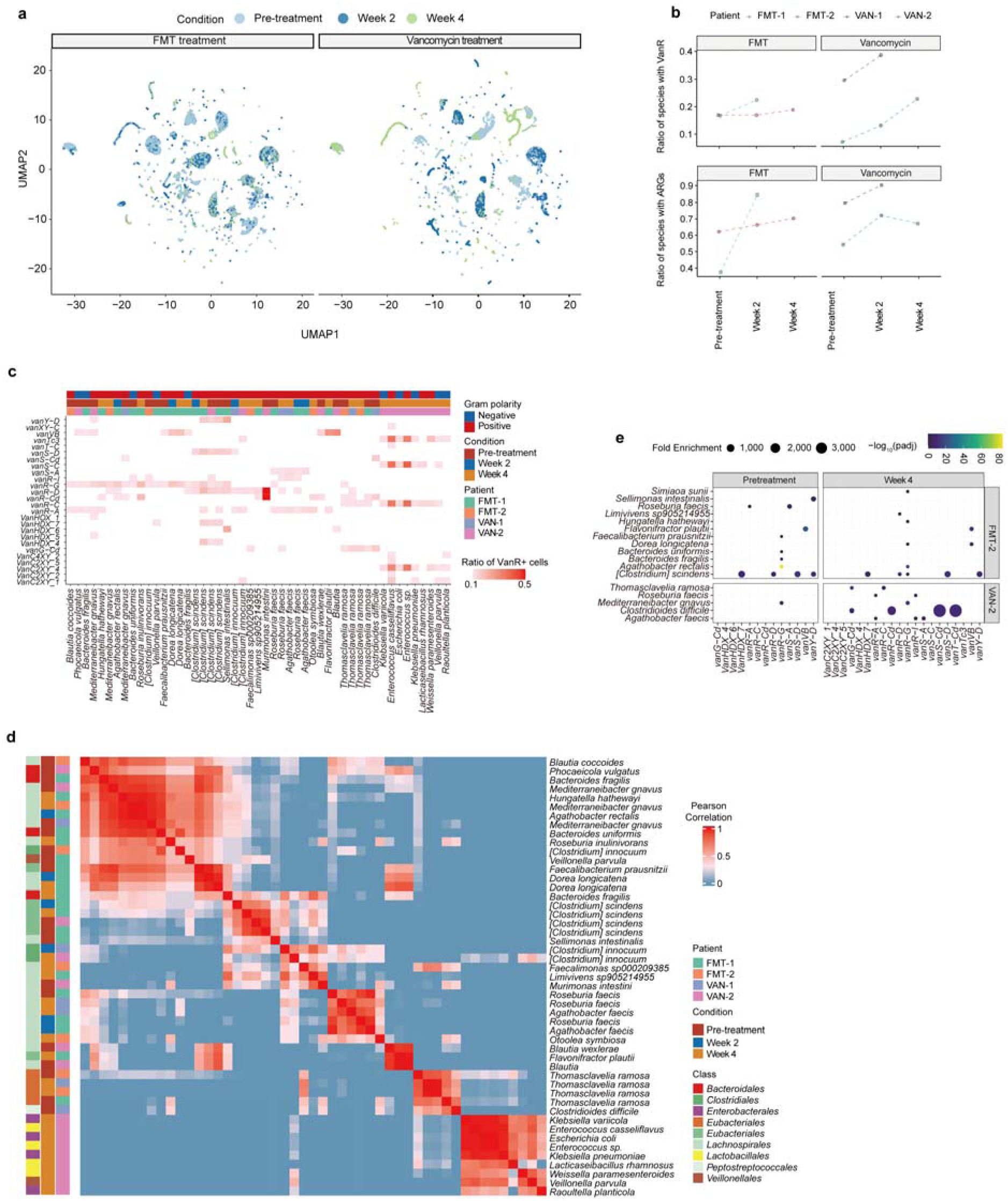
Single-cell profiling of treatment-induced shifts in microbial communities and resistance gene dynamics. (a) UMAP projections of SAGs from FTM- and vancomycin-treated patients across pre-treatment, week 2, and week 4. Each dot represents one SAG, colored by sampling stage. (b) Temporal changes in the proportion of species carrying vancomycin resistance genes (VanR, top) and total antimicrobial resistance genes (ARGs, bottom). (c) Taxonomic distribution of VanR genes across patients and time points. (d) Correlation matrix of VanR gene profiles across species. (e) Relative enrichment of VanR genes within host species across patients and treatment stages. Bubble size indicates fold enrichment, and color denotes statistical significance (-log_10_(adjusted P value)). Enrichment of multiple VanR genes in *C. difficile* was observed in patient VAN-2.

To identify which taxa drove these compositional changes, we compared species-level abundance shifts across consecutive treatment intervals and clustered taxa by their temporal trajectories (Supplementary Figs. 5b&c). Across both therapeutic regimens, only a limited subset of species exhibited pronounced abundance changes (> 0.05 in relative abundance), indicating that community restructuring was dominated by a few key responders. Notably, most taxa showing significant increases after vancomycin treatment were Gram-negative, whereas many Gram-positive commensals declined, consistent with the antibiotic’s narrow activity spectrum targeting peptidoglycan synthesis in Gram-positive bacteria. Among the expanding taxa, *Veillonella parvula* increased in both FMT- and vancomycin-treated patients. *B. fragilis* expanded after FMT but declined following vancomycin exposure. Given that both *Veillonella* and *Bacteroides* are not direct targets of vancomycin, their distinct abundance shifts may be the consequence of intervention-induced ecological restructuring. It can also be noted that the Gram-positive *Enterococcus casseliflavus* increased under vancomycin treatment, in line with its known intrinsic vancomycin resistance^28^. Together, these observations point to treatment-specific restructuring of gut communities, where FMT maintains higher continuity with baseline composition and vancomycin drives more pronounced turnover toward antibiotic-tolerant taxa.

Having characterized the taxonomic restructuring of gut communities under the two treatment regimens, we stepped to explore how these ecological shifts were accompanied by changes in the antimicrobial resistome. Specifically, we examined the dynamics of antimicrobial resistance genes (ARGs) during FMT and vancomycin treatment in CDI patients. Focusing on vancomycin resistance genes (VanR), we observed an increase in the proportion of species carrying VanR after treatment across all patients, with a more pronounced expansion in those receiving vancomycin (Fig. 5b, top). This stronger enrichment likely reflects the direct selective pressure imposed by vancomycin exposure, which favors the survival and expansion of intrinsically resistant taxa as well as the maintenance or horizontal transfer of VanR determinants among tolerant populations. When considering broader ARGs, a similar upward trend was evident in both FMT patients (Fig. 5b, bottom), indicating that FMT can also promote the dissemination of resistance genes within recipient microbiomes. These observations align with previous reports that donor-derived microbiota may introduce novel resistance determinants, underscoring the need to monitor ARG dynamics under both therapeutic strategies.

We then examined the taxonomic distribution of VanR genes across patients and treatment stages (Fig. 5c). Note that Fig. 5c only highlights taxa showing clear interspecies co-occurrence of VanR modules rather than all detected carriers. In VAN-2 at week 4, a distinct pattern emerged in which multiple taxa carried related VanR modules. Among these, the intrinsically resistant Enterococcus displayed the highest prevalence of these determinants, while several coexisting Gram-negative species, including Klebsiella pneumoniae, *E. coli*, *V. parvula*, and *Raoultella planticola*, harbored similar gene combinations. This co-occurrence suggests potential horizontal dissemination or shared maintenance of VanR modules across phylogenetically diverse hosts, possibly originating from *Enterococcus* under vancomycin selection. The presence of these resistance determinants in Gram-negative taxa may also explain their pronounced abundance increases during vancomycin therapy, as shown in Supplementary Figs. 5b and 5c, reflecting selective advantages conferred by acquired tolerance. Together, these observations reveal a cross-phyla resistance signature at the late stage of vancomycin therapy, where VanR modules spread among both diversified members of the perturbed gut community.

To further dissect the relationships among bacterial hosts carrying VanR genes, we next focused on the interspecies organization of resistance profiles. Fig. 5d captures taxon–taxon correlations based on shared VanR gene repertoires. Two major clusters with conserved resistance patterns emerged. The first cluster (upper left of the matrix) was dominated by members of *Lachnospirales* and *Bacteroidales* that were consistently detected across treatment stages and patients, suggesting a stable ecological module that maintains vancomycin resistance within commensal communities. The second large cluster (lower right corner) encompassed *Enterococcus* and several co-occurring taxa that shared correlated VanR signatures, corresponding to the cross-phyla resistance pattern observed in Fig. 5c for patient VAN-2 at week 4. In addition to these broad clusters, several smaller species-specific clusters were evident. For example, multiple *[Clostridium] scindens* and *Roseburia faecis* lineages formed tight blocks along the diagonal. These taxa displayed highly conserved resistance profiles that persisted across patients and time points, indicating stable maintenance of intrinsic or vertically transmitted VanR determinants. Together, these structured co-occurrence patterns reveal both large-scale ecological modules and lineage-specific conservation of resistance within the CDI gut microbiome.

Building on the species-level patterns, we further examined the organization of VanR genes themselves. Correlation analysis identified several compact clusters of co-occurring resistance determinants (Supplementary Fig. 5d), indicating structured relationships among VanR modules rather than random distribution. These associations likely reflect conserved genomic linkage or shared mobilization contexts, suggesting that VanR genes are frequently organized into coordinated cassettes that can be jointly maintained or transferred across microbial hosts.

At last, we explored how the VanR genes were associated with their bacterial hosts under different therapeutic contexts. Fig. 5e and Supplementary Fig. 5e display the relative enrichment (fold enrichment scores) of VanR genes within their hosts across patients and treatment stages. A notable enrichment was observed in patient VAN-2, where several VanR genes showed pronounced enrichment within C. difficile, potentially reflecting the early emergence or selection of a vancomycin-tolerant subpopulation within this species. Taken together, the above results illustrate the dynamic restructuring of the gut microbiome and its resistome during FMT and vancomycin therapy, revealing both treatment-specific and patient-specific trajectories that shape microbial composition, resistance potential, and ecological stability.

### Single-cell tracking reveals dynamic responses of pBI143 to therapeutic perturbations

Having characterized the temporal dynamics of microbial composition and resistance genes, we extended our analysis to plasmid dynamics, using pBI143 as a model to trace how mobile genetic elements and their bacterial hosts adapt during treatment. Overall, plasmid carriage ratio decreased progressively across most patients, during both FMT and vancomycin treatment (Fig. 6a), in line with broader ecological restructuring of the gut microbiome. This trend was consistent across treatment groups, although the rate of decline varied between patients. To resolve lineage-level dynamics, we tracked individual pBI143 versions across time (Supplementary Fig. 6). An overall decline in pBI143-carrying cells was observed. Among the four pBI143 versions, V1 remained the most prevalent at later stages. In patient VAN-2, nearly all detectable plasmids were lost by week 4, with only a few SAGs retaining V1, indicating a near-complete collapse of plasmid carriage under vancomycin-associated perturbation.

**Figure 6.**
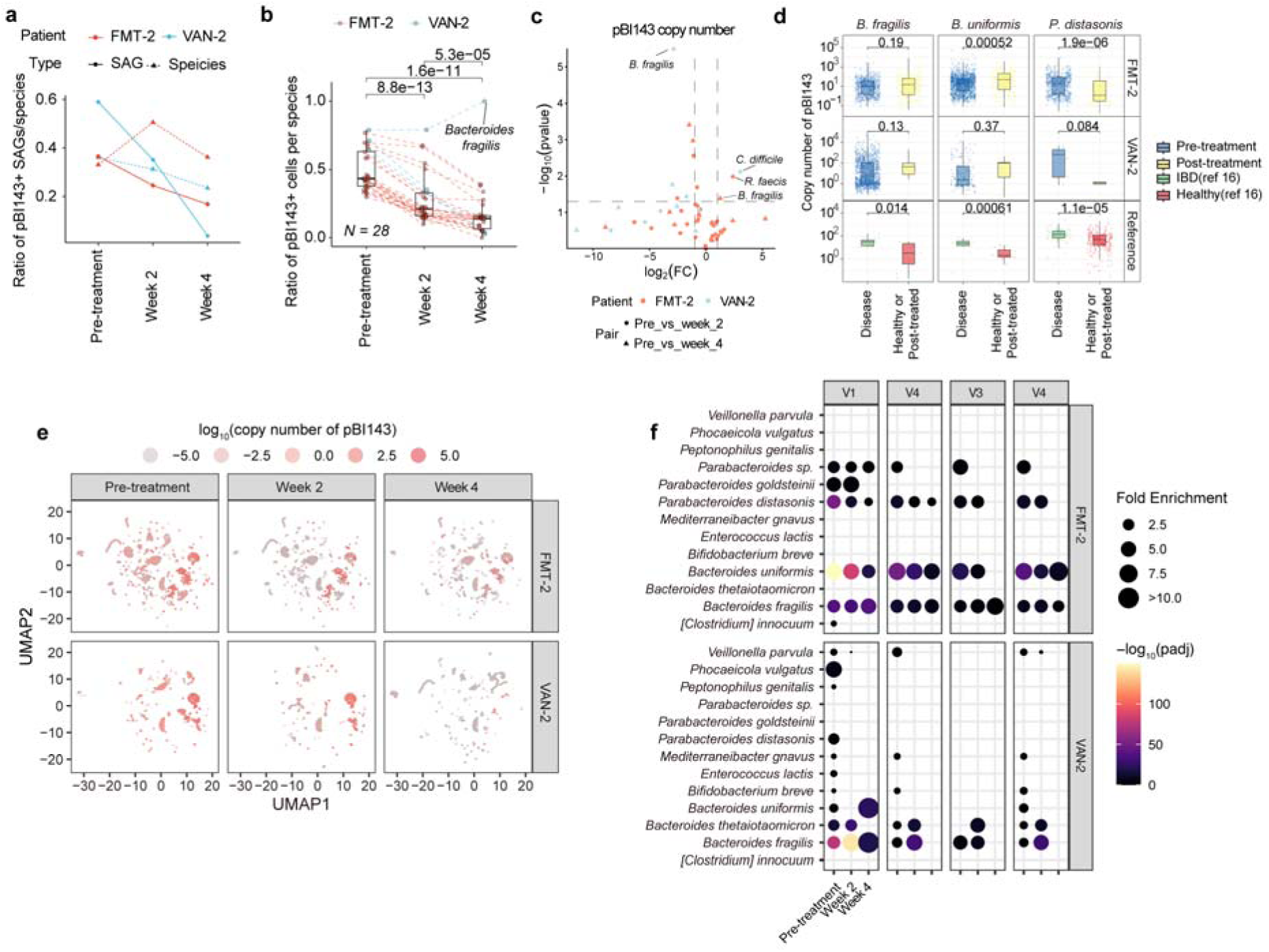
Single-cell tracking of plasmid pBI143 dynamics and host associations during therapeutic intervention. (a) Changes in the ratio of pBI143-carrying SAGs (circles) and host species (triangles) across treatment stages for patients FMT-2 and VAN-2. (b) Per-species ratio of pBI143-positive cells across stages for 28 recurrent taxa (carrying pBI143 in ≥1 sample). The specified numbers were P values obtained from paired two-tailed t-tests. (c) Differential analysis of pBI143 copy numbers across stages. The statistical significance was assessed by two-tailed t-tests. (d) Comparison of pBI143 copy number distributions in B. fragilis, B. uniformis, and P. distasonis among disease, post-treatment, and healthy reference cohorts (ref 16). The specified P values were obtained from two-tailed t-tests. (e) UMAP projections of gut microbiomes of patients FMT-2 and VAN-2 at different treatment stages, showing changes in pBI143 copy numbers. (f) Fold enrichment of pBI143 versions (V1–V4) across host species and treatment stages. Bubble size denotes fold enrichment and color indicates statistical significance (–log₁₀ padj).

We then identified which bacterial hosts contributed to these treatment-associated changes. Focusing on 28 species that appeared across all three sampling stages and carried the plasmid in at least one sample (Fig. 6b), most taxa exhibited marked decreases in the fraction of pBI143-positive cells after therapy, indicating a broad contraction of its ecological range. *B. fragilis* was the only notable exception, showing an increase in plasmid carriage, suggesting possible host-specific retention or fitness advantages.

We further examined plasmid copy number dynamics within these host species (Fig. 6c). Across most species, copy numbers remained stable or decreased after treatment. Notably, *C. difficile* exhibited a significant increase in plasmid copy number in patient VAN-2, suggesting that this taxon may provide a permissive niche for plasmid replication during treatment. Interestingly, while *B. fragilis* showed an overall increase in the fraction of plasmid-carrying cells (Fig. 6b), its copy number dynamics were heterogeneous across patients. This discrepancy may suggest a decoupling between population-level plasmid spread and intracellular plasmid amplification. Such patterns may arise from host strain–specific plasmid replication control or ecological pressures that differentially shape plasmid maintenance strategies across individuals. Together, these observations highlight that pBI143 copy number dynamics during the therapy are species- and patient-specific, rather than uniform across the community.

To place these findings in context, we compared pBI143 copy-number distributions in *B. fragilis*, *B. uniformis*, and *P. distasonis*, three species overlapping with the prior metagenomic study^16^ of pBI143 (Fig. 6d). In our CDI cohort, *P. distasonis* exhibited a significant elevation in plasmid copy number during disease, which decreased following treatment, consistent with previous observations^16^. For *B. fragilis* and *B. uniformis*, copy-number differences trended lower in disease compared to post-treatment and healthy reference samples, but most differences were not statistically significant. These results indicate that disease-associated plasmid amplification may be species-specific rather than a universal feature across hosts.

To visualize plasmid copy number dynamics at the single-cell level, we mapped pBI143 abundance onto UMAP projections of all SAGs across treatment stages (Fig. 6e). High-copy clusters (red) were prominent before treatment but progressively diminished during both FMT and vancomycin therapy, consistent with overall plasmid loss (Figs. 6a-c). However, several high-copy clusters persisted at week 4 for both patients. To capture such dynamics in pBI143 host preference throughout the treatment, we assessed the fold enrichment of individual pBI143 versions across treatment stages (Fig. 6f). At the pre-treatment stage, all four plasmid versions were broadly distributed across diverse taxa, showing moderate associations with multiple *Bacteroides*, *Parabacteroides*, and *Veillonella* species. As treatment progressed, these associations became increasingly focused, with enrichment signals converging toward the *Bacteroides* lineage, particularly *B. fragilis* and *B. uniformis*. This progressive narrowing of host range suggests that the therapeutic perturbation selectively favors plasmid persistence in *Bacteroides* while depleting its presence in other taxa.

Together, these results demonstrate that pBI143 responds dynamically to therapeutic perturbation at multiple ecological scales. While overall prevalence and copy number decline, certain host–plasmid combinations can persist, underscoring the context-dependent resilience of mobile genetic elements during microbial community restructuring.

## DISCUSSIONS

CAP-seq combines the strengths of high-throughput single-cell sequencing with a streamlined, microfluidics-minimal workflow, overcoming key limitations of existing methods. By performing critical reactions within permeable hydrogel compartments, CAP-seq enables efficient cell lysis, whole-genome amplification, and library preparation in bulk, thereby improving reaction yield and genome recovery while substantially simplifying the overall process. In contrast to current high-throughput single-cell genomics methods, which typically rely on multiple serial microfluidic operations, CAP-seq requires only two (cell encapsulation and barcode pairing), thereby substantially reducing experimental complexity. This design makes the method highly scalable and amenable to automation, similar in spirit to commercial droplet-based transcriptomic platforms such as 10x Genomics^29^. Moreover, the workflow can be further simplified by replacing the encapsulation step with bulk shaking emulsification^30^, or by adopting split-pool barcoding strategies^31^ for genome indexing, potentially enabling a fully microfluidics-free implementation. These features collectively position CAP-seq as a practical and flexible platform for large-scale single-cell genomic studies across diverse environments.

Our results demonstrate that CAP-seq bridges a critical gap between metagenomics and single-cell genomics, enabling scalable, high-resolution exploration of microbial communities at unprecedented depth. By capturing thousands of individual microbial genomes in parallel, CAP-seq makes it possible to directly link genetic elements, such as plasmids, phages, and resistance determinants, to their cellular hosts and ecological contexts. This level of resolution provides new opportunities to investigate microbial population structure, horizontal gene transfer, and genome evolution within natural ecosystems and host-associated microbiomes. Looking forward, further improvements in amplification fidelity, read length, and integration with complementary single-cell modalities (e.g., transcriptomics) could transform CAP-seq into a comprehensive platform for multi-omic microbial cell profiling. Much like the impact of single-cell RNA-seq on eukaryotic biology, scalable microbial single-cell genomics promises to reshape our understanding of microbial ecology, evolution, and their roles in health and disease.

## METHODS

### Microfluidics

Microfluidic devices were designed using AutoCAD (v2023) and fabricated using polydimethylsiloxane (PDMS)-based soft lithography. Before use, the devices were treated with Aquapel (PPG Industries) to render the channel surfaces hydrophobic. The microfluidic experiments were conducted on a custom-built microfluidic station consisting of an inverted microscope equipped with a high-speed camera (Phantom VEO310L) and six syringe pumps (New Era NE-510). The flow rates of all inlets in the experiments were specified in Supplementary Figs. 1a&b.

### Generation and characterization of CAPs

The CAPs were generated using a flow-focusing microfluidic chip (Supplementary Fig. 1a and Supplementary Movie 1) by emulsifying an aqueous two-phase system (ATPS) consisting of dextran and acrylate-modified poly(ethylene glycol) diacrylate (PEGDA) in the continuous phase of fluorinated oil (3M HFE-7500) with 2% v/v biocompatible polyethylene glycol-perfluoro polyether (PEG-PFPE) surfactant (DP-Bio FSA2). To prepare the ATPS solutions, a premix was prepared in a 1.5 mL centrifuge tube. A 1 mL premix contained: 1x DPBS buffer, Dextran (M.W. 500,000; Yeason 61216ES25; 0.55 g), PEGDA (M.W. 8,000; Alfa Aesar 046801.03; 0.03 g), PEGDA (M.W. 575; Sigma-Aldrich 437441; 30 μL), 4% (w/v) LAP (Lithium phenyl-2,4,6-trimethylbenzoylphosphinate; Sigma-Aldrich 900889) (100 μL). After mixing, the tube was centrifuged at 14,000 rpm for 30 min, resulting in a clear separation between an upper PEGDA-rich phase and a lower dextran-rich phase. The two phases were then collected separately for subsequent emulsification. Upon emulsification, phase separation within the droplets resulted in a capsular configuration, with PEGDA forming the outer layer. The CAPs were collected in a tube and irradiated under a UV lamp (200 Watt) for 4 min to gel the outer PEGDA phase. The semi-permeability of the CAPs was characterized using a customized PCR within the CAPs, where we amplified DNA fragments (RhaT gene) of varying lengths from encapsulated *E. coli* genomic DNA. This experiment determined the threshold size of dsDNA products that can be trapped in the CAPs. Detailed procedures for the characterization can be found in our previous reports^12,13^. The CAPs containing intact and lysed cells, and amplified genomes (Scheme 1) were stained by SYBR Green (Biotium 40086) and examined under a fluorescent microscope.

### Cell culture

a. *E. coli* (strains BL21 and MACH1), *B. subtilis* (strain ster164), *S. aureus* (strain RN4220), and *K. pneumoniae* (strain KP_1.6366) were separately cultured in 5 mL of LB medium at 37°C overnight in a shaking incubator. The cell concentration was determined by manually counting serial dilutions of the liquid culture on plastic slides (ThermoFisher C10228) under a microscope.

### Stool samples

Stool samples were collected at the initial diagnosis of CDI, before treatment with antibiotics (vancomycin) or FMT, and at weeks 2 and 4 afterward. All samples were collected at the Department of Gastroenterology, Hepatology and Nutrition, Shanghai Children’s Hospital, and stored at 4 °C for immediate usage. For long-term storage, the samples were mixed with 25% v/v glycerol and stored at -80°C.

To extract the microbial cells from stool, the samples were resuspended in PBS containing 0.1% Pluronic F68 (ThermoFisher, 24040032) by vigorous vortexing. Large debris was then removed by centrifugation at 4°C at 35g for 20 minutes. The cell-containing supernatant was transferred to a new tube, washed twice with PBS + 1% Pluronic F68, and resuspended in 100 µL of PBS + 1% Pluronic F68. Cell concentration was evaluted by manual counting.

### CAP-seq procedure

#### Cell encapsulation

Cell samples were resuspended in PBS buffer and spiked in the Dextran-rich solution (the inner phase of CAP) to a final cell density of 600 CFU/μL. This led to a positive rate of ∼5% to minimize doublet formation, according to the Poisson statistics. The CAPs were then generated using microfluidics and collected in a 1.5 mL centrifugal tube. Upon gelling by UV irradiation, the CAPs were extracted from the oil by adding an equal volume of 20% Perfluoroctane (PFO; Sigma-Aldrich 359238) in fluorinated oil and washed three times with DPBS buffer containing 0.1% v/v Tween-20 (Sigma-Aldrich P6585).

#### Cell lysis

To lyse the encapsulated cells, 500 μL of dense-packed CAPs were mixed with an equal volume of 20 mM TE, 100 mM NaCl, 10 mM EDTA, 1% Triton X-100 (ThermoFisher 85111), 1800 U lysozyme (Lucigen R1804M) and 20 U lysostaphin (Sigma-Aldrich L7386). The mixture was incubated at 37 °C for 2 hours. The CAPs were then washed with PBS buffer containing 10 mM EDTA for three times and mixed with 500 μL of 100 mM NaCl, 10 mM EDTA, 1% SDS, and 1mg proteinase K (ThermoFisher AM2546). The mix was incubated at 55°C for 30 min. The CAPs were eventually washed with a washing buffer containing 10 mM HEPES, 2% Tween 20, 20 mM EDTA and 0.1 mM NaCl for three times and resuspended in a DPBS buffer containing 0.1% Tween-20.

#### Whole genome amplification

After lysis, 300 μL of dense-packed CAPs were mixed with an equal volume of MDA mix containing 2X Phi29 reaction buffer (Vazyme N106- 02), 30 μL Phi29 polymerase (Vazyme N106-02), 60 μL Pyrophosphatase (ThermoFisher EF0221), 20 mM dNTP (Vazyme P031-01), 0.25% v/v Triton X-100 and 0.25% v/v Pluronic F68. The MDA reaction was incubated at 30°C for 2 hours and terminated at 65°C (10 min). The post-MDA CAPs were washed with the washing buffer three times and resuspended in a DPBS buffer containing 0.1% Tween-20.

#### Tagmentation

200 μL of the post-MDA CAPs were mixed with an equal volume of tagmentation solution containing 80 μL Tag buffer (ABclonal RK20239) and 40 μL Tag Enzyme 50 (ABclonal RK20239; aliquot with nuclease-free water). The mix was incubated at 55°C for 10 min. Afterward, the CAPs were washed with the washing buffer three times and resuspended in a DPBS buffer containing 0.1% Tween-20.

#### Genome Barcoding

To barcode the tagmented genomes, the CAPs were co-encapsulated with barcode beads and a linear amplification mix using a custom bead- pairing device^20^ (Supplementary Fig. 1b). The linear amplification mix (total volume 348 μL) consisted of 120 μL Phusion High-Fidelity Reaction Buffer (NEB M0530L), 16 μL Phusion High-Fidelity DNA Polymerase (NEB M0530L), 16 μL Deep Vent DNA Polymerase (NEB M0258L), 12 μL ET-SSB (NEB M2401S), 64 μL 100 mM DTT (Sigma-

Aldrich D9779), 64 μL 10× PCR stabilizer (DP-Bio RRF0981), 64 μL 10 mM dNTP, and 16 μL nuclease-free water. The barcode bead (DTT-dissolvable polyacrylamide hydrogel bead) carried ssDNA primers containing a unique barcode sequence and an adapter sequence designed to bind to the genome fragments. The beads used in this study were custom-ordered from PercentBio, China (RRF0991), and the fabrication process has been described elsewhere^32^. The applied flow rates in the barcoding step are detailed in Supplementary Fig. 1b. After merging, the resulting droplet contained a CAP and a unique barcoding primer (a dissolved bead), and the linear amplification mix.

For linear amplification, the droplets were collected in PCR tubes and thermal cycled with the following recipe: 72°C for 5 min, 98°C for 3 min, and 15 cycles of 98°C for 10s, 60°C for 30s and 72°C for 5 min. Before thermal cycling, the original emulsification oil was replaced by Bio-Rad QX200 oil.

#### CAP dissolution

After barcoding, the droplets were broken by adding an equal volume of 20% PFO in HFE-7500. The released CAPs were washed three times using TE buffer containing 0.1% Tween-20. Next, the CAPs were then dissolved by mixing 20 μL dense-packed CAPs (containing ∼9,000 cells) and 40 μL 0.8 M NaOH and incubated at 55°C for 10 min. Subsequently, the resulting solution was neutralized by adding 40 μL 0.8 M acetic acid. The DNA products were then purified by magnetic beads (ThermoFisher AMPure XP) and dissolved in 15 μL nuclease-free water.

#### Bulk PCR and sequencing

The barcoded genomic DNA library was further amplified by PCR. The PCR reaction contained 15 μL purified linear amplification product, 25 μL KAPA hot-start HiFi polymerase mix (Roche 07958935001), and 5 μL each of the forward and reverse primers. The PCR condition was 98°C for 3 min, and 16 cycles of 98°C for 10s, 60°C for 30s and 72°C for 5 min. The PCR product was purified using magnetic beads, examined using a Bioanalyzer (Agilent 2100), and then used for Nanopore sequencing library preparation (using Ligation Sequencing Kit V14, SQK- LSK114, and Native Barcoding Kit 24 V14, SQK-NBD114.24) following the manufacturer’s protocol (https://nanoporetech.com/products/prepare/dna-library-preparation). The final DNA library was sequenced on an Oxford Nanopore sequencer (GridION MK1 with P2 solo) using a PromethION Flow Cell (FLO-MIN114). All the primer sequences for CAP-seq are detailed in Supplementary Table 1.

### Bioinformatics

#### Initial quality control

Raw sequencing data in POD5 format were base-called using Dorado (v0.8.3) in sup mode. The resulting reads were filtered with NanoFilt (v2.8.0) to remove low-quality sequences (quality score < 15) and reads with lengths outside the range of 20 to 60,000 bases.

#### Barcode identification and demultiplexing

Cell barcodes were identified by locating a known adapter sequence (GTCTCGTGGGCTCGG) using the Python (v3.14) pairwise2.align.localms function from the Bio module, with alignment parameters set as pairwise2.align.localms(seq1, seq2, 2, -1, -1, -1). Matches with an alignment score of 26 or higher were considered successful, and the adapter position was used to further locate the associated barcode sequence. Extracted barcodes were validated against a predefined whitelist of expected sequences. Reads with valid barcodes and complete library structures were annotated by appending the barcode sequence to their header. Following barcode annotation, adapter sequences were removed using Cutadapt (v5.1). Demultiplexing was then performed with BBMap’s (v39.37) demuxbyname.sh script, generating individual FASTQ files, each corresponding to a single SAG. Codes for barcode identification and demultiplexing are archived on GitHub (https://github.com/Liulab2023/CAP-seq).

#### Purity and coverage assessment

Purity and genome coverage metrics were calculated to evaluate the quality of each valid SAG in the sequencing of the *E. coli* and *B. subtilis* mixture and mock community. Purity was determined as the proportion of reads within a SAG mapped to the dominant species, based on alignment results from Minimap2 (v2.30). Genome coverage was quantified using Samtools (v1.22.1) by calculating the percentage of the reference genome covered by aligned reads. For the *de novo* assemblies of the mock microbial sample (Fig. 2e), genome completeness and contamination were evaluated with CheckM (v1.0.13).

#### *De novo* assembly

SAGs were assembled using Miniasm (v r179) with the following parameters: miniasm -i 0.03 -m 50 -s 50 -e 1 -g 2000 -F 0.5 -c 1 -n 0 -1 -2. Assembly quality was evaluated with QUAST (v5.3.0).

#### Microbiome species annotation

Assembled genomes were aligned against the RefSeq, GTDB databases, and a curated library of human gut bacterial isolates (PRJNA544527). Species-level classification was based on Average Nucleotide Identity (ANI ≥ 95%).

#### UMAP analysis

To visualize single-cell genomic heterogeneity, we performed UMAP dimensionality reduction on pairwise ANI distances between SAG contigs and their reference genomes (n = 1,702). Pairwise ANI was computed using skani (v0.3.0) with thresholds for ANI and alignment fraction (AF) set to 0 to retain all distance information, and -c 30 to accommodate short fragments. SAG pairs with alignment fractions <15% were filtered to reduce noise. UMAP was then applied using cosine similarity, producing the single-cell atlas shown in Fig. 3b.

#### Phylogenetic tree construction

Phylogenetic trees for both reference species (Fig. 3c) and SAGs (Fig. 3f) were constructed from pairwise ANI distances computed with skani (thresholds set to 0, -c 30). In Fig. 3c, we selected the top 100 most abundant species across samples. For species mapped to multiple genomes, the maximum ANI value across versions was used. Distance matrices were generated using cosine similarity, and trees were built by complete-linkage hierarchical clustering with hclust in R.

#### SAG SNP and coverage calculation

SNPs and InDels were called by aligning SAG contigs to their reference genomes using snippy (v4.6.0) with default settings (Figs. 3c&g). For coverage estimation, SAG contigs were aligned using minimap2, and per-site coverage was calculated with samtools. The same approach was used for pBI143 coverage (Fig. 4a).

#### Identification and assembly of pBI143

Reads from each SAG were aligned to the pBI143 reference (GenBank U30316.1). SAGs with >98% plasmid coverage were retained for plasmid assembly using Rebaler (v0.2.0).

#### pBI143 version identification and sequence construction

To characterize plasmid diversity, contigs covering the full length of the pBI143 reference were collected (coverage > 0.98; n = 5,247). High-quality contigs (a contig constructed by >100 reads in its original SAG; n = 2,010) were selected, aligned with MAFFT (v7.525), and clustered using K80 distances (ape::dist.dna) and complete linkage (hclust). Four clusters (versions) were defined by cutree, and consensus sequences were generated with consensusString. Versions were assigned to remaining contigs based on maximum AF and ANI using skani (v0.3.0), excluding ambiguous ties. In total, 5,177/5,247 contigs were assigned to one of four plasmid versions. Full sequences of the four identified versions of pBI143 were deposited on the National Genomics Data Center of China (Project No. PRJCA047574).

#### Analysis of pBI143 copy number dynamics

To analyze plasmid copy number changes during the treatments of CDI, we selected species with ≥5 plasmid-positive cells in all samples per individual. Post-treatment samples were stratified by time point (2 and 4 weeks). For each species and patient, per-cell copy number was averaged pre- and post-treatment, and log₂ fold change was computed. Significance was determined using a two-tailed t-test (p < 0.05 and |log₂FC| > 1). For comparisons with previous studies (Fig. 6d), week 2 and week 4 samples were merged into a single post-treatment group.

#### pBI143 Version Enrichment Analysis

To determine whether specific versions of the pBI143 plasmid were enriched within particular bacterial species, a Fisher’s exact test-based enrichment analysis was performed. For each unique combination of pBI143 version, bacterial species, experimental group, and treatment condition, a 2×2 contingency table was constructed using the following calculated counts:

a: The number of unique SAGs containing the target pBI143 version within the specific bacterial species.

b: The number of unique SAGs of the specific species that did not harbor the target pBI143 version (calculated as the total number of cells of that species, cellnum_species, minus a).

c: The number of unique SAGs harboring the target pBI143 version in all other species (calculated as the total number of cells with this pBI143 version, cellnum_pBI143, minus a).

N: The total number of SAGs from all species within the respective experimental group and condition, serving as the background population. The value for d (SAGs not belonging to the target species and not harboring the target plasmid) was derived as (d = N - (a + b + c)).

A one-sided Fisher’s exact test (alternative hypothesis: "greater") was applied to each contingency table to test the null hypothesis that the pBI143 version was not enriched within the target species. The odds ratio (OR) was calculated to quantify the strength of association. The fold enrichment was computed as ((a/(a+b))/(c/(c+d))) to provide an intuitive measure of the enrichment magnitude. To ensure statistical robustness, analyses were only performed for combinations where the value of a was greater than 5. A P-value of less than 0.05 was considered statistically significant.

#### Jaccard Similarity Analysis

To assess the similarity in bacterial species composition between samples, we employed Jaccard similarity coefficient analysis (Supplementary Fig. 5a). First, a presence-absence matrix was constructed from the input data, where rows represent individual samples and columns denote unique bacterial species. Each entry in the matrix indicates whether a given species is present (1) or absent (0) in a particular sample.

The Jaccard similarity coefficient was calculated using the formula:

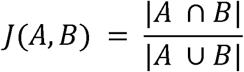

where A and B are sets of species present in two compared samples. Box plots (Supplementary Fig. 5a) were generated to compare Jaccard scores across treatment conditions, including statistical testing (paired t-test) to evaluate significant differences in microbial community shifts induced by treatments such as FMT versus Vancomycin.

#### Analysis of Temporal Changes in microbial species

The differences in cell ratios were calculated for each species by subtracting the baseline value (pre-treatment) from the values at subsequent time points (week 2 and week 4). Then we filtered out species with minimal overall change (sum of absolute differences ≤ 0.05) to focus on biologically relevant variations. Box plots were created to detail the magnitude and distribution of cell ratio changes for individual species (Supplementary Fig. 5b). Additionally, a heatmap was generated to display the filtered difference matrix, with columns ordered by experimental group and time point to facilitate the comparison of temporal trends across treatments (Supplementary Fig. 5c).

#### Identification of VanR and ARGs

To obtain the distribution of *VanR* and broader ARGs on the assembled SAGs, we used the Abricate tool (v1.0.1) to align the assembled genome against drug-resistance databases, including CARD, ARGANNOT, NCBI, and ResFinder. A resistance gene was considered if the alignment coverage was greater than or equal to 10% and the sequence identity was greater than or equal to 70%.

#### Enrichment Analysis of VanR and ARGs

The enrichment of *VanR* and other ARGs within distinct bacterial species was assessed using a methodology analogous to the pBI143 enrichment analysis. For each unique combination of target ARG, bacterial species, sample, and experimental group, the core parameters (a, b, c, d, N) for the contingency table were calculated as described in the pBI143 analysis. A one-sided Fisher’s exact test (alternative hypothesis: "greater") was applied to each ARG-species combination to test the null hypothesis of no enrichment. The odds ratio and fold enrichment were calculated. To avoid spurious results from low counts, the analysis required SAG counts to be greater than 2. A P-value of less than 0.05 was considered statistically significant. This method allows for the direct assessment of ARG enrichment within specific bacterial populations from complex metagenomic data.

#### Correlation Analysis of Antimicrobial Resistance Genes and Species Distribution

To investigate the co-occurrence patterns of ARGs across different bacterial species and experimental conditions, a correlation analysis followed by hierarchical clustering visualization was performed. The initial dataset contained binary presence/absence records for each ARG within each bacterial species, stratified by experimental group and condition. The data were first subsetted to exclude unclassified or non-bacterial entries. A data matrix was constructed where each row represented a unique ARG, and each column represented a unique combination of species, group, and condition. The values represented the total occurrence count of each ARG within each specific context. Columns with a sum of occurrences less than or equal to 1 were filtered out to ensure robust correlation calculation.

A correlation matrix was computed from the transposed data matrix using Pearson correlation coefficients, which quantified the pairwise similarity in ARG presence/absence patterns across all contexts. To focus on the most interconnected associations, the matrix was filtered to retain only rows (i.e., species-group-condition combinations) where the row sum of correlation coefficients exceeded a threshold of 1.2, isolating contexts with substantial co-occurrence relationships.

## DATA AVAILABILITY

All data supporting the findings of this study are available within the article and its Supplementary Information. The raw sequencing data of this study are archived on the National Genomics Data Center of China (https://ngdc.cncb.ac.cn) with project number PRJCA047574.

## CODE AVAILABILITY

All codes for the bioinformatics of this study are available at the following GitHub repository: https://github.com/Liulab2023/CAP-seq.

## AUTHOR CONTRIBUTIONS

(a) Y. Liu conceived the CAP-seq method. M. Zhang, J. Li, and R. Zhang developed and optimized the CAP-seq workflow under the supervision of Y. Liu, with assistance from J. Zhang. Z. Song contributed to CAP-seq library preparation. M. Li and Y. Du performed CAP-seq sequencing of mock and clinical samples. X. Li collected stool samples. S. Li, H. Liu, X. Zhai, and J. Tong established the bioinformatic pipeline. Y. Liu, H. Liu, S. Li, X. Zhai, J. Tong, and W. Wei carried out data analysis. Y. Liu, W. Wei, and T. Zhang supervised the microbiome study. Y. Liu, H. Liu, and S. Li prepared the figures and wrote the manuscript. Y. Liu, W. Wei, T. Zhang, Y. Luo, and Q. Ji secured funding for this work.

## ACKNOWLEDGEMENT

The authors thank Dr. Jian Li for kindly providing the *E. coli* and *B. subtilis* strains, and Drs. Lichun Jiang, Ruyuan Song, and Suwen Zhao for insightful discussions. Part of the bioinformatic analyses were performed on the High-Performance Computing Platform of ShanghaiTech University. This work was supported by the start-up funding of ShanghaiTech University, the National Natural Science Foundation of China (32300081), the Shanghai Science and Technology Committee (23QA1406600), the ShanghaiTech AI4S Initiative (SHTAI4S202404), and the Development Funds of the School of Physical Science and Technology (SPST-YSFZ-2024-01) and the School of Life Science at ShanghaiTech University.

## COMPETING INTERESTS

(a) Y. Liu, Y. Du, J. Li, and R. Zhang are co-inventors on a patent application filed by ShanghaiTech University based on the CAP-seq workflow described in this article. Y. Liu and W. Wei are co-founders of a startup company aimed at commercializing CAP-seq.

---

**Scheme 1.**
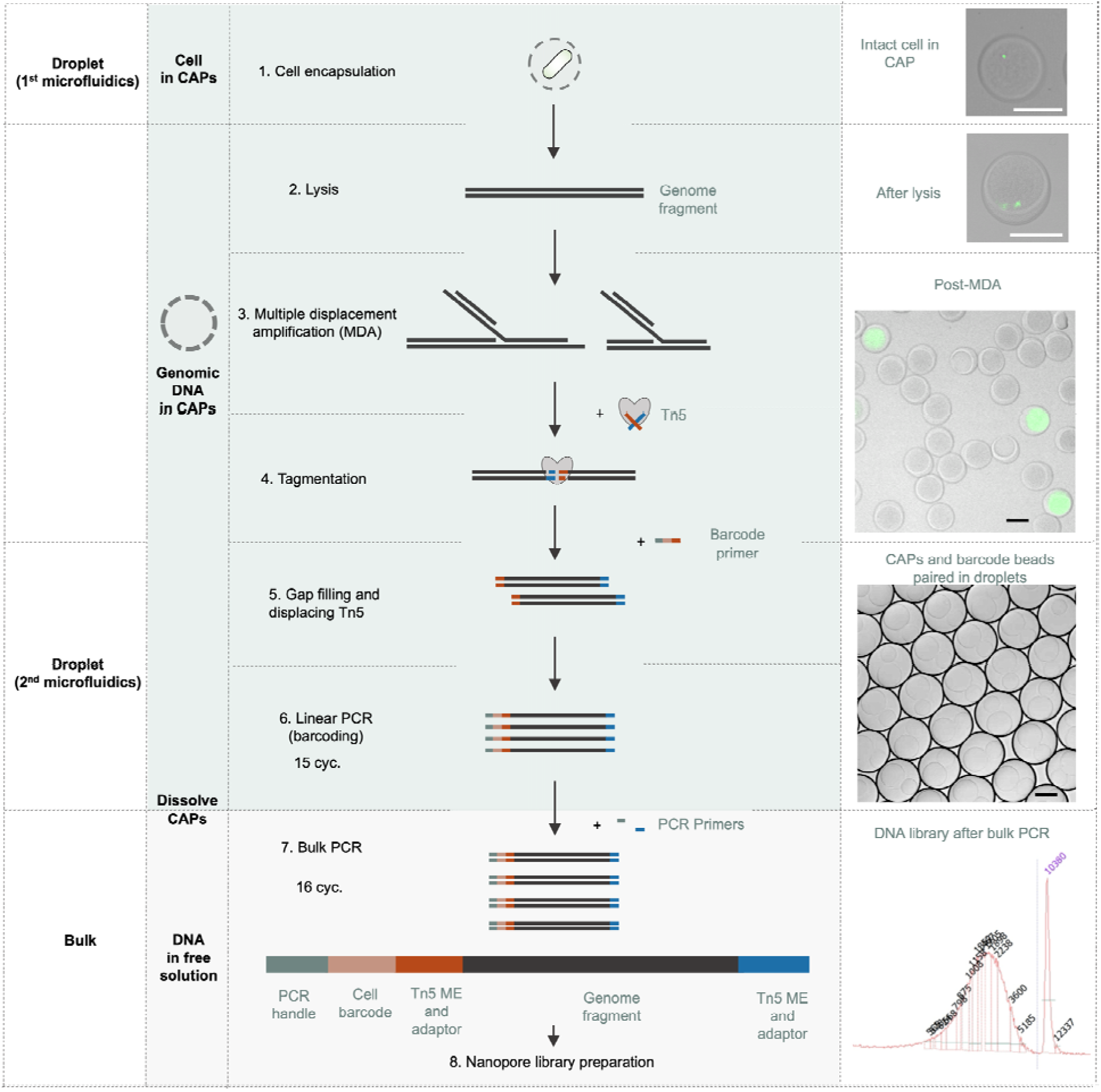
The CAP-seq workflow and key quality checking steps. Scale bars: 50 µm.

**Supplementary Figure 1.**
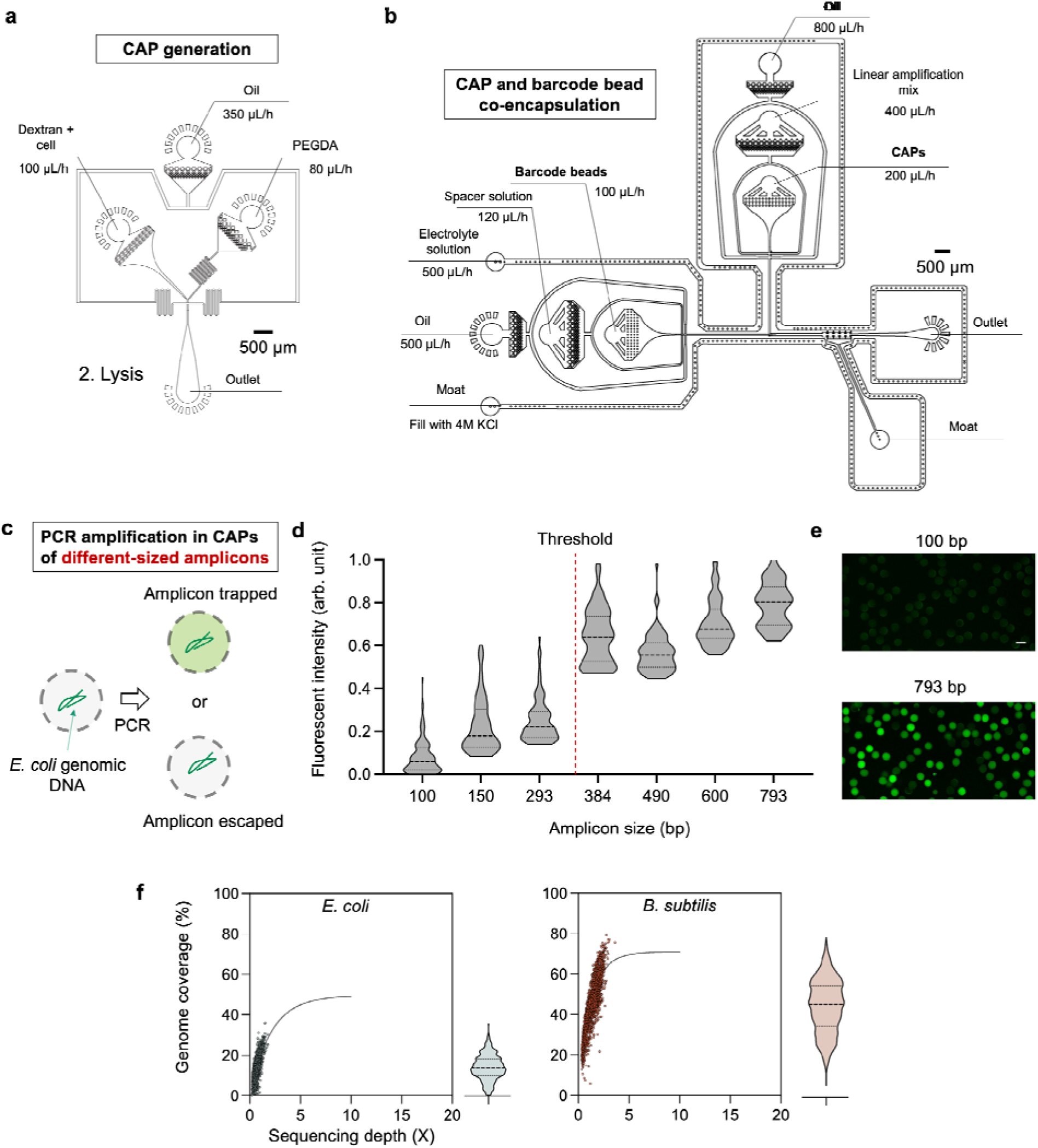
(a&b) Microfluidic designs and fluidic conditions of microfluidic experiments in CAP-seq: (a) CAP generation and (b) co-encapsulation of CAPs containing amplified SAGs and barcode beads. (c-e) PCR-based assay to assess the permeability of CAPs. (c) Schematic of the assay. *E. coli* genomic DNA templates were encapsulated in CAPs, and multiple PCR experiments were carried out, amplifying amplicons of varying sizes. Amplicons smaller than the pore size diffuse out of the capsule, whereas larger fragments remain trapped. (d) Quantification of fluorescence intensity of CAPs after PCR. Signal retention increased sharply for fragments larger than ∼400 bp, defining the effective cutoff for dsDNA (N > 70 for each amplification). (e) Representative fluorescence images showing size-dependent amplicon retention within CAPs (100 bp vs 793 bp). Scale bar: 50 µm. (f) Genome coverage versus sequencing depth for single *E. coli* (left; N = 1,105) and *B. subtilis* (right; N = 1,751) cells processed by CAP-seq. Each dot represents one barcoded SAG. The curved lines are rarefaction fitting curves showing the tendency of coverage saturation with increasing sequencing depth. The right panels summarize coverage distributions across all recovered SAGs.

**Supplementary Figure 2.**
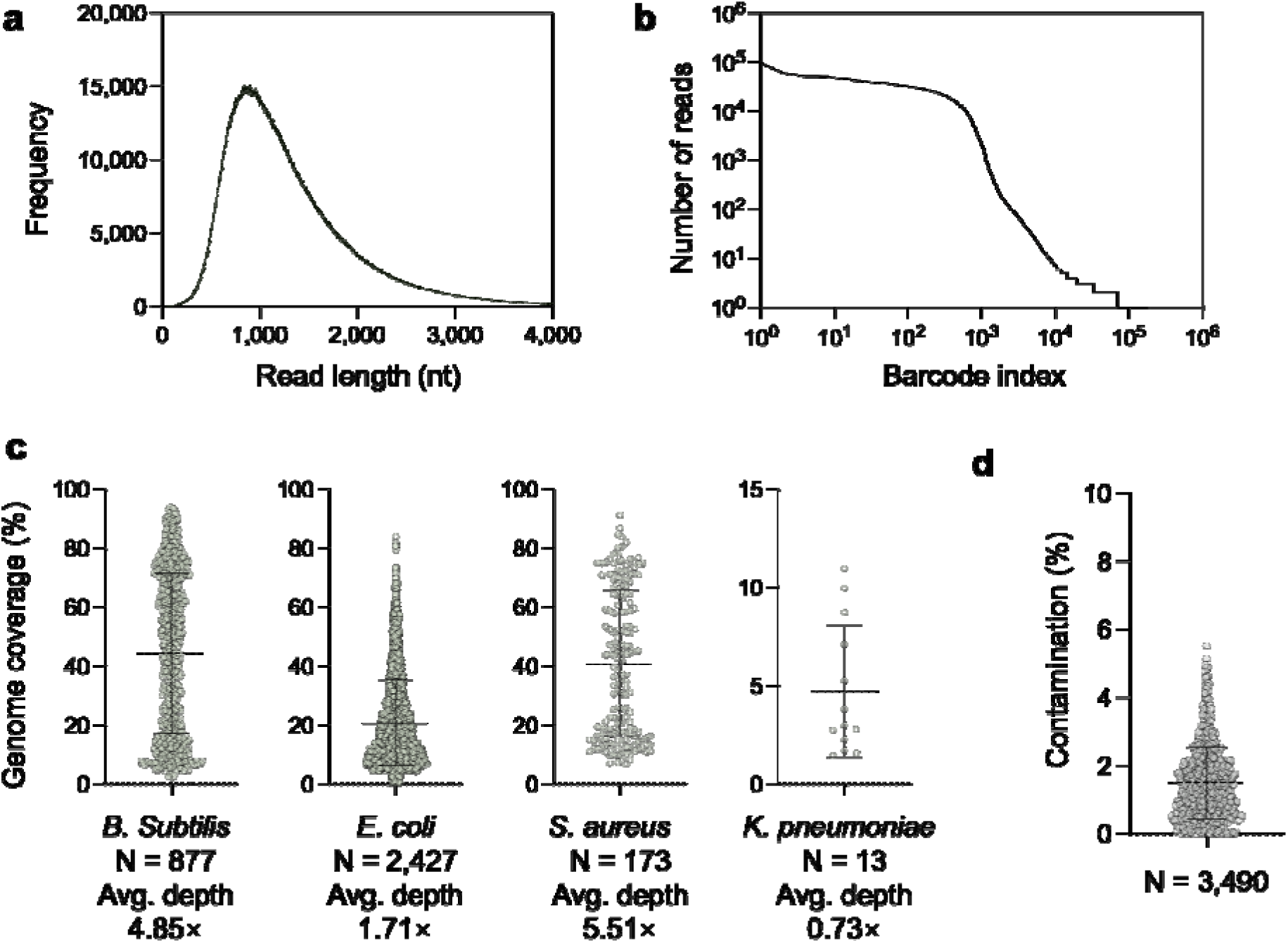
(a&b) Quality metrics of the CAP-seq experiment of Fig. 2. (a) Read length (after barcode trimming) profile. Bin size: 1. (b) Read distribution across different SAGs. (c) Scatter plots showing genome coverage as a function of sequencing depth for all SAGs. (d) Scatter plot showing the contamination levels of the *de novo* assemblies.

**Supplementary Figure 3.**
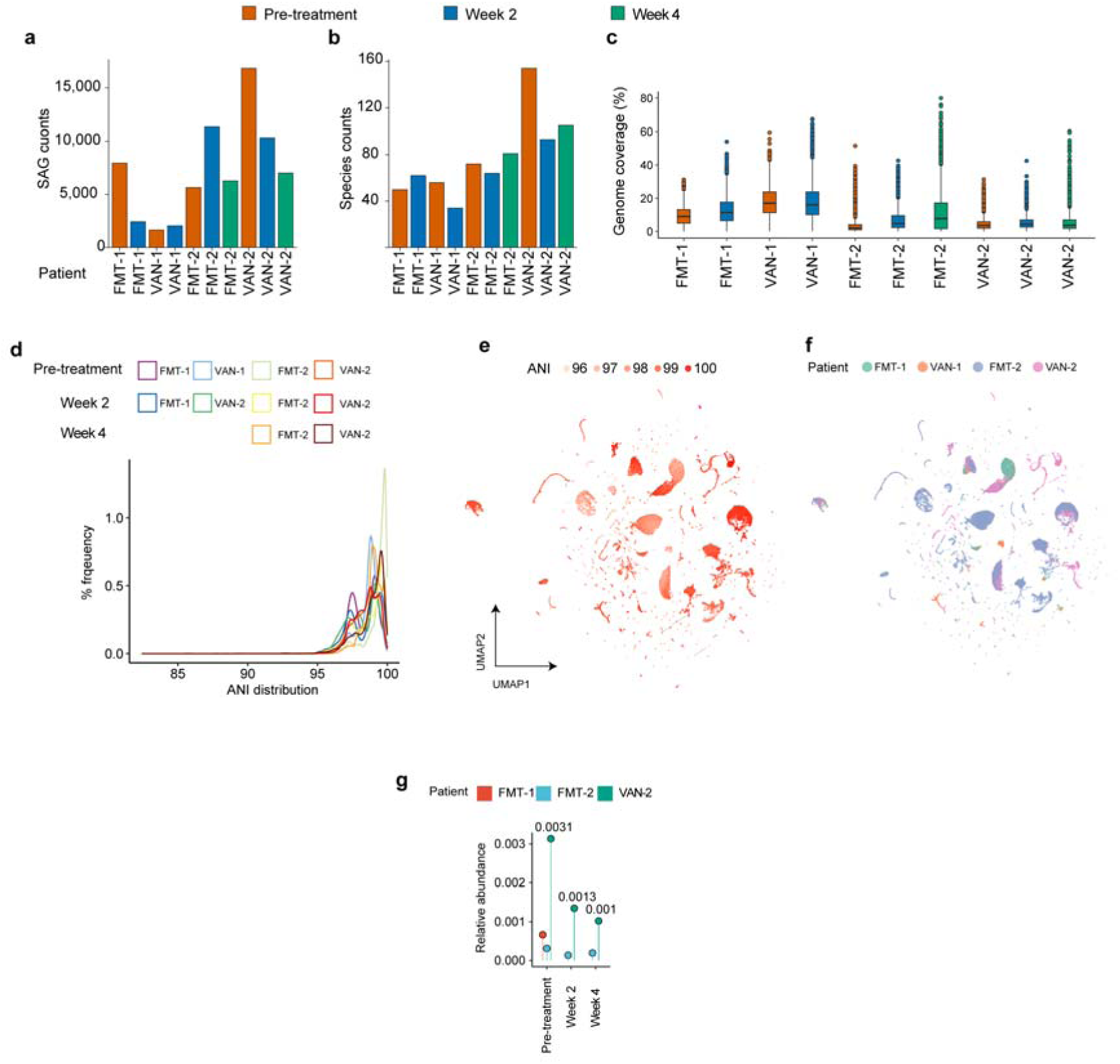
(a) Number of SAGs recovered from each of the four CDI patients across treatment stages. (b) Corresponding number of species identified per sample after *de novo* assembly and taxonomic annotation. (c) Distribution of genome coverage across all SAGs. (d) Distribution of average nucleotide identity (ANI) values among all SAGs. (e) UMAP projection of all SAGs colored by ANI values. (f) UMAP projection colored by patient identity. (g) Relative abundance of *C. difficile* across samples and treatment stages.

**Supplementary Figure 4.**
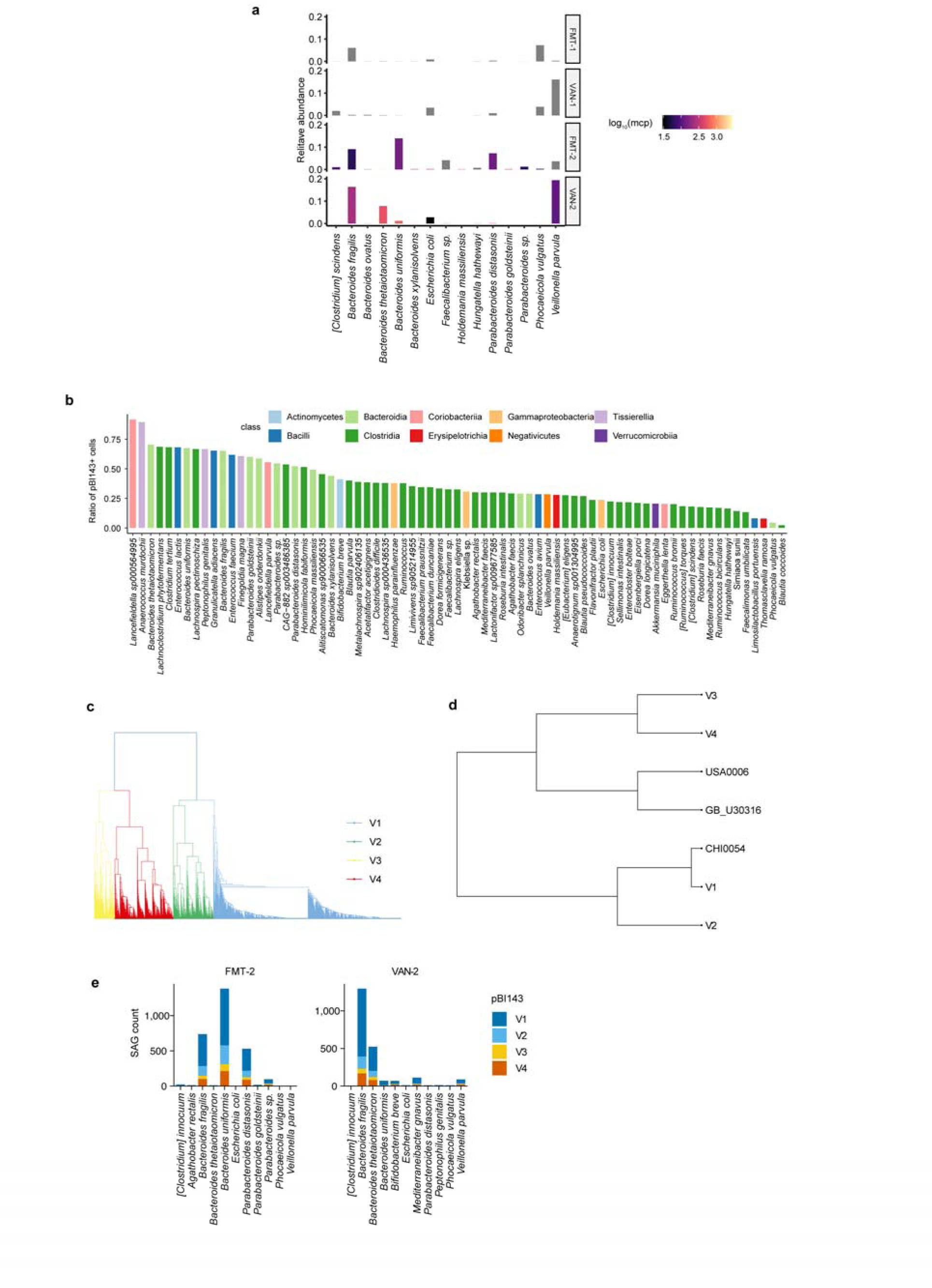
(a) Relative abundance of major bacterial species carrying pBI143 across all four CDI patients. (b) Fraction of pBI143-positive SAGs within each bacterial species, colored by taxonomic class. (c) Hierarchical clustering of pBI143 contigs based on SNP profiles, identifying four distinct plasmid versions (V1–V4) recovered by CAP-seq. (d) Phylogenetic relationships between the four versions (V1–V4) identified in this study and reference pBI143 lineages (CHI0054, USA0006, GB_U30316), showing partial relatedness but clear genetic divergence. (e) Distribution of plasmid versions across dominant host species in patients FMT-2 and VAN-2.

**Supplementary Figure 5.**
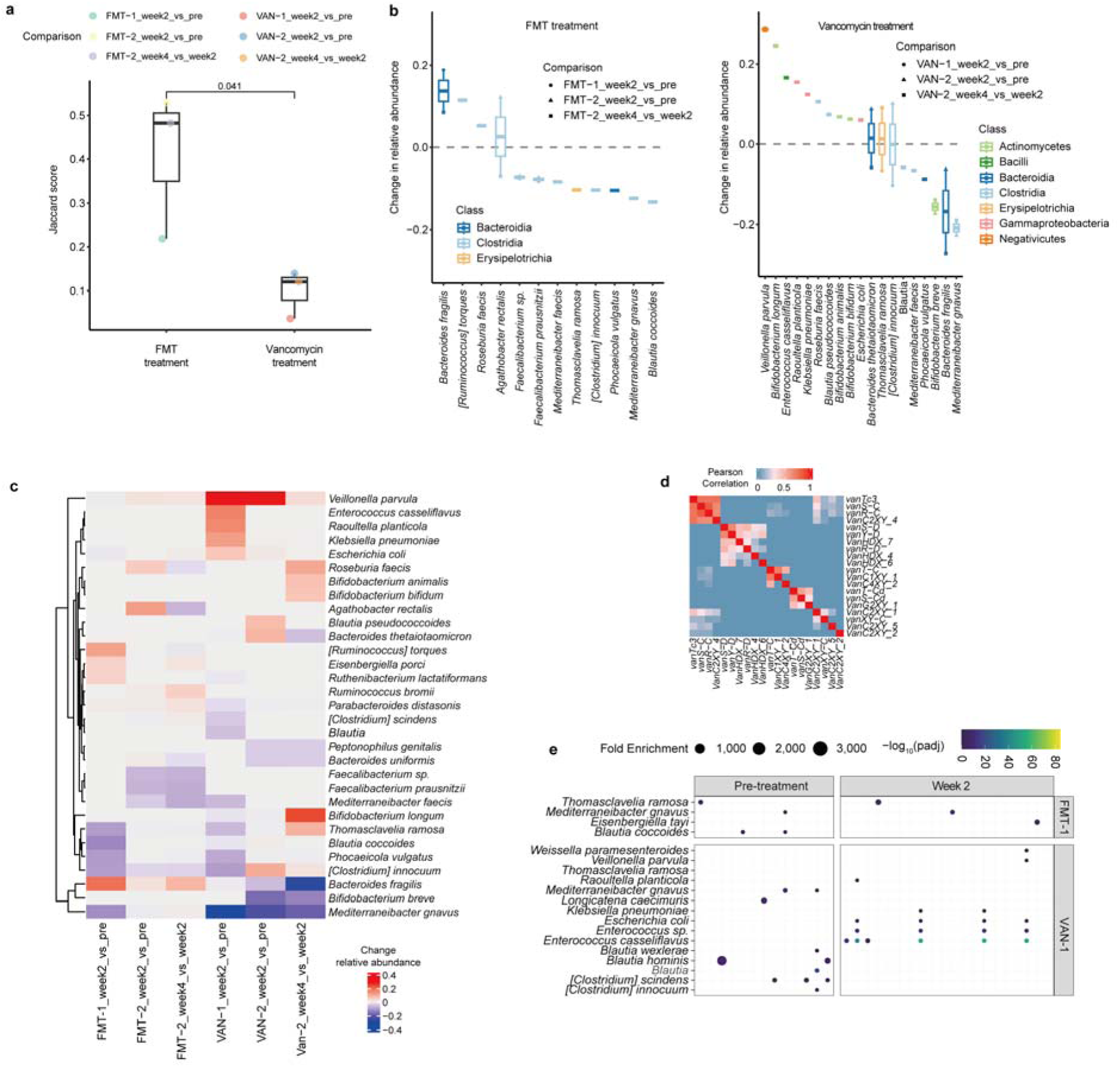
(a) Pairwise Jaccard similarity between consecutive sampling stages, quantifying compositional turnover during FMT and vancomycin treatment (P value = 0.041 by two-tailed t-test). (b) Species-level abundance changes between consecutive treatment stages for FMT (left) and vancomycin (right) patients. (c) Hierarchical clustering of species by temporal abundance changes, revealing therapy-specific restructuring modules. (d) Pearson correlation matrix of VanR gene profiles across bacterial species. (e) Fold enrichment of VanR genes within bacterial hosts across patients and treatment stages.

**Supplementary Figure 6.**
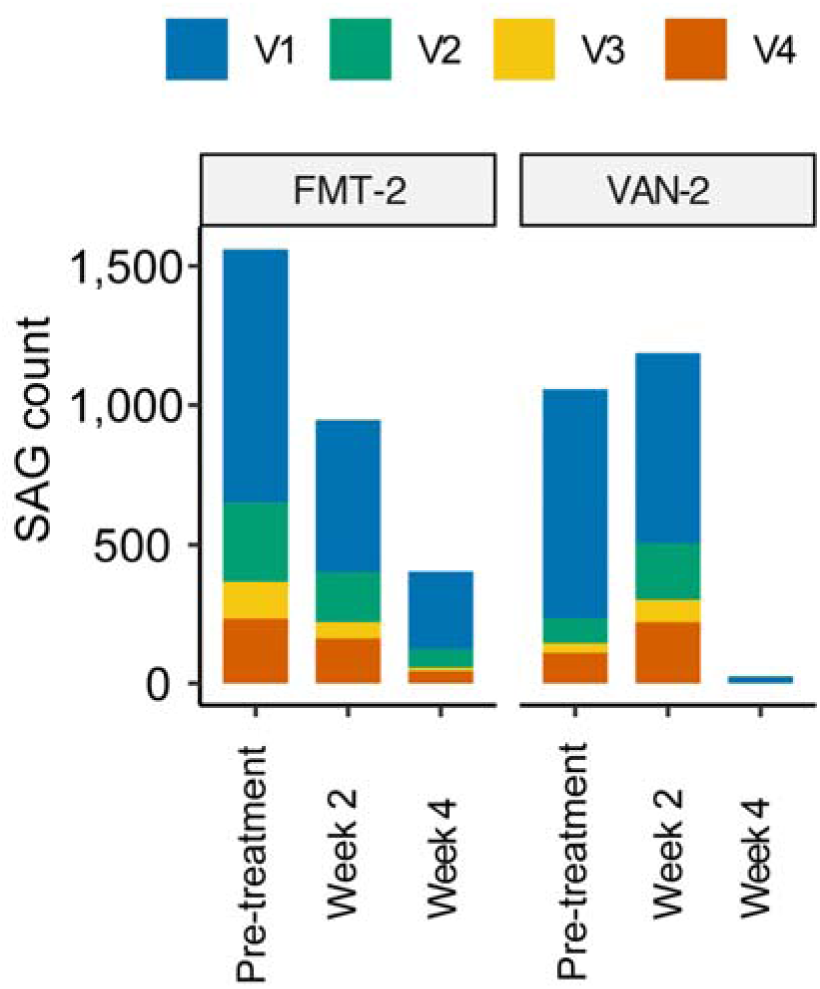
Stacked bar plots showing the abundance (plasmid-containing SAG counts) of four pBI143 plasmid versions (V1–V4) across treatment stages in two CDI patients (FMT-2 and VAN-2).

## Notes

### Competing Interest Statement

The authors have declared no competing interest.

### Summary of Updates

A new version of the manuscript

